# Establishment, maintenance and consequences of inter-individual transcriptional variability for a gene involved in nitrate nutrition in plants

**DOI:** 10.1101/2025.04.10.648086

**Authors:** Charlotte Lecuyer, Alexandre Vettor, Cécile Fizames, Hélène Javot, Antoine Martin, Mona Mazouzi, Sandra Cortijo

## Abstract

Differences in phenotypes and gene expression are observed between genetically identical plants grown in the same environment. While we now have a good knowledge of the source and consequences of transcriptional differences observed between cells, in particular in unicellular organisms, it is still very scarce when it comes to variability between multicellular organisms. Using *NRT2.1*, a high affinity nitrate transporter gene, as a model for high inter-individual transcriptional variability, we showed that differences in expression between plants for that gene are established in young seedlings, maintained over time but not transmitted to the next generation. Our results also indicate that these differences in expression could have phenotypic consequences. Indeed, they can explain phenotypic variability for the root growth as well as the amount of nitrate imported by the NRT2.1 protein in the plant. Finally, we found an enrichment for genes involved in photosynthesis among the ones with an expression correlated with *NRT2.1* in single seedlings. All in all, our study suggests that a global coordination of the genes involved in the carbon/nitrate balance in plants is established in young seedlings, with differences in this state between plants, and then maintained over time.

## Introduction

Genetically identical plants grown in the same environmental conditions can display differences in phenotypes and gene expression. Interestingly, this inter-individual variability has been observed in plants for seed germination timing (Abley et al., 2021; Mitchell et al., 2017), aerial phenotypes like phyllotaxy (Kitazawa, 2021), the plant height, rosette size or number of flowers (Hall et al., 2007) as well as lateral root length (Forde, 2009). The level of this phenotypic variability is at least partially genetically controlled, as shown by several successful GWAS and QTL studies in which they identified regions of the genome that can influence the level of inter-individual variability in the different genotypes (Jimenez-Gomez et al., 2011; Shen et al., 2012; Abley et al., 2021; Hall et al., 2007; Joseph et al., 2015). Moreover, it has also been shown that transcriptional inter-individual variability is widespread, with 9% of the transcriptome displaying a high level of variability (Cortijo et al., 2019). The existence of this variability can have both positive and negative consequences. On one hand, it has been proposed that variability in seed germination could help populations of plants in the desert to better survive in case of unexpected drought after a rainfall (Cohen, 1968; Mitchell et al., 2017; Childs et al., 2010; Abley et al., 2024). Moreover, there are several examples in unicellular organisms where the variability helps populations of cells to survive a stress that would be otherwise lethal (Grimbergen et al., 2015; Morawska et al., 2022). On the other hand, variability for a range of phenotypes like seed germination or flowering time are not necessarily desirable for crops, for which a strong inter-individual homogeneity is expected. Another aspect to keep in mind is that the individual (in a population) is the level at which selection happens during evolution. In this context, it is essential to understand where the transcriptional variability is coming from in plants as well as its phenotypic consequences.

While there is now a good number of studies of transcriptional noise observed between cells, the knowledge of inter-individual variability of gene expression in multicellular organisms is still scarce. It is known that the source of cell-to-cell transcriptional noise is the inherent stochasticity of the transcriptional process itself, with layers of factors that can either enhance or buffer this stochasticity (Cortijo and Locke, 2020; Dong and Liu, 2017). These factors can come from the genomic, epigenomic or regulatory circuit structures as well as the cell age, global transcriptional state of the cell or its microenvironment (Tantale et al., 2016; Sharon et al., 2014; Tirosh and Barkai, 2008; Dey et al., 2015). When it comes to the differences in expression observed between multicellular organisms, knowing the source of this variability is more complex. Indeed, since they are composed of thousands of cells, it is not clear if the factors that are known to be involved in the variability between cells are also the one explaining the variability between multicellular organisms. A transcriptomic study of inter-individual variability in *Arabidopsis thaliana* showed that genes that are highly variable have a genomic and epigenomic structure similar to the one observed in unicellular organisms (Cortijo et al., 2019), suggesting the same mechanisms are at play. However, we are missing a lot of information to understand where the inter-individual variability is coming from in plants, as well as whether it can lead to phenotypic consequences. For example, we do not know when inter-individual variability is established during the life of plants, if it responds to the environment or if differences in expression between plants are maintained over time.

To answer these questions, we decided to follow the variability over time and in different environmental conditions for one highly variable gene, using imaging approaches. We chose the *NRT2.1* gene as a model for several reasons. First of all it has all the characteristics of a highly variable gene from our previous study: it is an environmentally responsive gene (Bellegarde et al., 2017; Medici and Krouk, 2014; Ruffel et al., 2019), is regulated by a high number of transcription factors (Bellegarde et al., 2017) and has a compact chromatin environment (Bellegarde et al., 2018). Moreover, this gene is a high affinity nitrate transporter, and we found that a high proportion of genes involved in nitrate nutrition are highly variable in our previous work (Cortijo et al., 2019). The presence of high variability for many genes in this pathway is of interest when we know that the presence, and concentration, of nitrate in the soil fluctuates spatially and temporally in natural conditions and fields (Cain et al., 1999; Shahandeh et al., 2005; Lin et al., 2019). Finally, *NRT2.1* is an important nitrate transporter, especially in conditions with low concentrations of nitrate in the soil where it is responsible for approximately 75% of the nitrate taken by the roots (Cerezo et al., 2001; Filleur et al., 2001). All of this makes *NRT2.*1 a good model gene to study the impact and establishment of inter-individual transcriptional variability.

In this study, we show that *NRT2.1* has a high inter-individual transcriptional variability in the environmental conditions we tested, as long as the gene is expressed. Most importantly, this variability in expression for *NRT2.1* is correlated with differences between seedlings for several phenotypes such as the root growth and the activity of the NRT2.1 protein in importing nitrate. We found that differences in expression between plants for that gene are established early on, shortly after the heterotrophic-to-autotrophic transition, are maintained for at least several days but not transmitted to the next generation. Finally, we observed that genes co-expressed with *NRT2.1* are not involved in nitrate nutrition but in photosynthesis highlighting a regulation for that gene centred around the nitrate/carbon balance in plants.

## Results

### The expression for the nitrate transporter *NRT2.1* is highly variable between plants

To define the level of plant-to-plant transcriptional variability for *NRT2.1* we performed RT-qPCR on RNA extracted from individual seedlings grown on a media containing a low concentration of nitrate. In these conditions, *NRT2.1* is known to be highly expressed and its function is essential to plant’s growth (Cerezo et al., 2001; Filleur et al., 2001). By doing so on multiple seedlings, we then measured the Coefficient of Variation (CV) that is the standard deviation divided by the average expression level in the seedlings. The highest the CV, the highest the variability for the gene analysed. We found that *NRT2.1* has a CV equivalent to a gene known to be highly variable (HVG, AT1G08930) and higher than for a gene with a low level of plant-to-plant transcriptional variability (LVG, AT2G28810) (Figure 1A). These two control genes come from a previously published study of the inter-individual variability for the entire transcriptome of *Arabidopsis thaliana* (Cortijo et al., 2019). This result shows that *NRT2.1* expression is indeed highly variable between plants.

**Fig 1.**
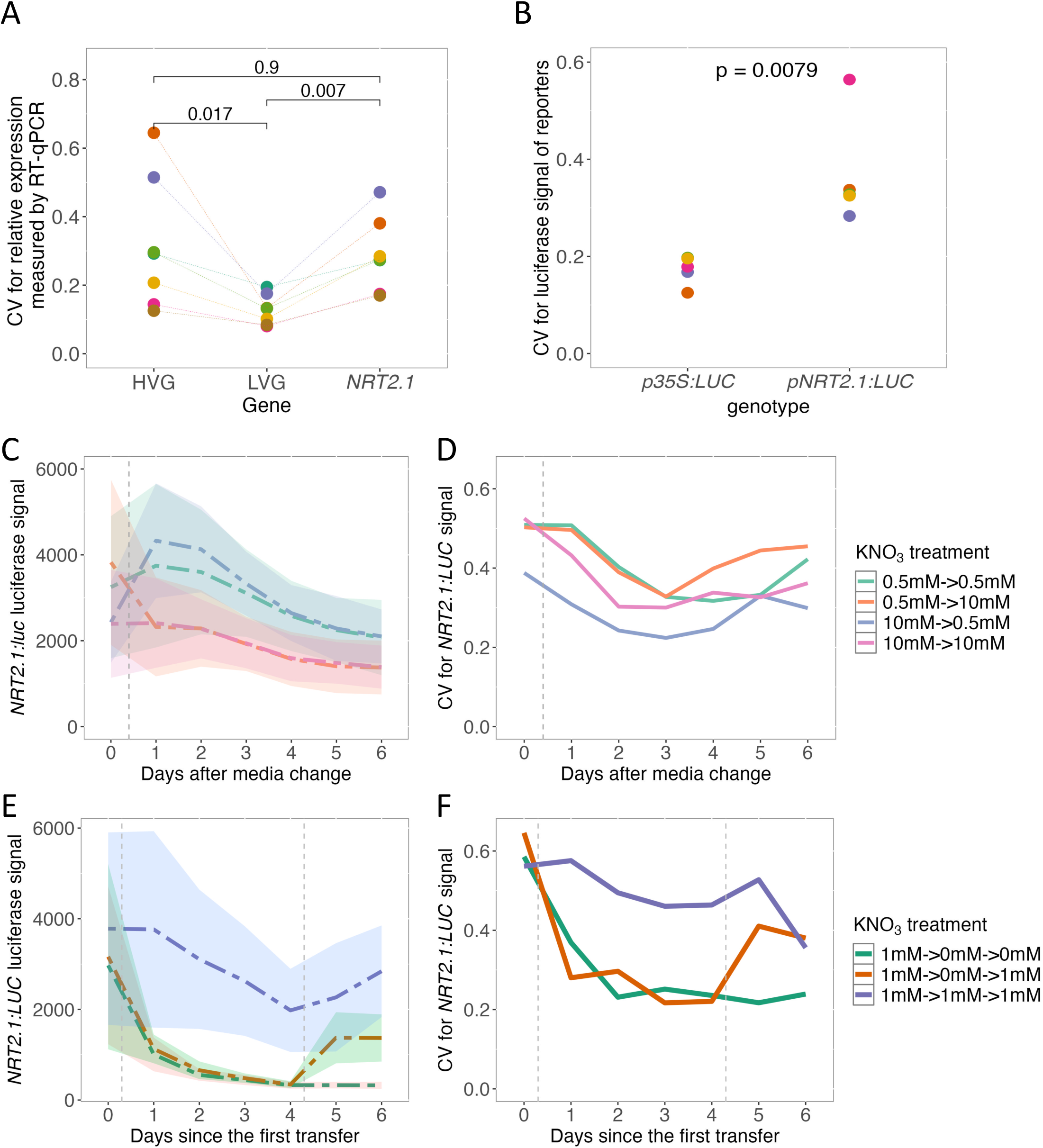
*NRT2.1* expression is highly variable between plants. (A) Inter-individual transcriptional variability measured by RT-qPCR for a highly variable gene (HVG), a lowly variable gene (LVG) and *NRT2.1*. The variability is measured using the CV (standard deviation/mean). Each colour represents a replicate with 24 seedlings per replicate grown on a media with low nitrate concentration (1mM). The results of a Wilcoxon test are included. (B) Measurement of inter-individual variability for *pNRT2.1* and *p35S* promoters using the *pNRT2.1:LUC and p35S:LUC* luciferase reporter lines respectively. Each colour represents a replicate with at least 28 seedlings per replicate grown on a media with low nitrate concentration (1mM). The result of a Wilcoxon test is included. (C-D) Signal (C) and CV (D) for the *pNRT2.1:LUC* reporter line in response to changes in nitrate concentrations. Dotted lines represent the mean signal (C) and solid lines the CV (D) at the different time points of the experiment measured on around 30 seedlings per condition. The vertical grey dotted line indicates the transfer of seedlings to a new plate. (E-F) Signal (E) and CV (F) for the *pNRT2.1:LUC* reporter line in response to nitrate starvation. Dotted lines represent the mean signal (E) and solid lines the CV (F) at the different time points of the experiment measured on around 30 seedlings per condition. The vertical grey dotted lines indicate the transfer of seedlings to a new plate.

We wanted to confirm this result by other means thanks to a promoter reporter line. For this, we used a luciferase promoter reporter line for *NRT2.1* that was already described (Girin et al., 2010) to measure the level of variability between plants for the promoter activity and compared it with a *p35S:LUC* line (Dorbe et al., 1998). The cauliflower mosaic virus (CaMV) 35S promoter allows for a constitutive and high expression of the luciferase reporter, making it a good control for low inter-individual variability. We found a higher inter-individual variability for the luminescence signal of *NRT2.1* promoter compared to the *35S* promoter (Figure 1B). This result confirms *NRT2.1* high variability between plants, whatever the method used to measure it. It also shows that the *pNRT2.1:LUC* reporter line can be used to study *NRT2.1* variability. Indeed, with luciferase imaging being faster and cheaper than RT-qPCR, we decided to use this approach to analyse more in detail the inter-individual transcriptional variability for *NRT2.1*. To further check if we can use this approach, we tested by RT-qPCR if *NRT2.1* endogenous gene and the luciferase gene under the control of the *NRT2.1* promoter are well co-expressed in single seedlings. We observed a strong positive correlation between the two (Figure S1A) indicating they are well co-regulated in single seedlings. The *pNRT2.1:LUC* reporter line can thus be used as a proxy for the *NRT2.1* endogenous gene.

### *NRT2.1* stays highly variable with different nitrate concentrations

All the analyses above were done under low nitrate conditions where *NRT2.1* is highly expressed. However it is known that *NRT2.1* expression level as well as its function is influenced by the concentration of nitrate on which the plants are growing (Nazoa et al., 2003; Medici and Krouk, 2014; Bellegarde et al., 2017). We wanted to test if the high inter-individual transcriptional variability for *NRT2.1* can also be affected by this. We thus analysed the variability for seedlings that were grown with either low (1mM KNO_3_) or high (10mM KNO_3_) nitrate concentrations using the *pNRT2.1:LUC* reporter line. It is important to keep in mind that none of these two conditions (low or high nitrate) are perceived as a stress for the plant as they stay in a physiological range. As already published (Nazoa et al., 2003; Medici and Krouk, 2014; Bellegarde et al., 2017), when combining 23 experiments, with around 25 seedling growing on low and 25 seedling growing on high nitrate in each experiment, we found that *NRT2.1* expression is higher for the plants grown with a low nitrate concentration (Figure S1B, 1.7 times on average). When it comes to the variability, we observed no statistically significant differences for the CV of plants grown with low or high nitrate concentrations (Figure S1C). To go further, we also analysed the impact of a fluctuation in the nitrate concentration on *NRT2.1* expression and variability by transferring plants growing on a low nitrate concentration in a new plate with a high nitrate concentration and vice-versa. To enhance the transcriptional response to this environmental change, we used bigger differences in nitrate concentration with 0.5mM and 10mM of KNO_3_ as low and high concentration respectively and this specifically for this experiment. We see that *NRT2.1* expression is behaving as expected: it increases for plants transferred to a media with a lower concentration of nitrate and goes down when plants are transferred to a media with a higher concentration of nitrate (Figure 1C). Again, we found that *NRT2.1* inter-individual transcriptional variability is not affected by the fluctuation in nitrate concentration in the media, whether plants are experiencing an increase or reduction of nitrate concentration (Figure 1D). These results suggest an uncoupling of the regulation of gene expression level and inter-individual transcriptional variability in response to nitrate concentration in the environment, since in both experiments gene expression of *NRT2.1* is affected by environmental changes when its variability is not.

We then decided to analyse a more severe change in the environment by studying the impact of a complete nitrate starvation on *NRT2.1* expression and inter-individual variability. For this, seedlings were grown on low nitrate conditions (1mM KNO_3_) for 6 days (T0) and then transferred on a new plate without any source of nitrogen. In parallel, controls were transferred on a plate with low nitrate. After four days of this treatment (10-day-old seedlings, T4), half of the starved seedlings were transferred again for one day to a plate with low nitrate, while the others were maintained under complete nitrate starvation (Figure S1D for experimental set-up). We observed that *NRT2.1* expression is gradually lost under complete nitrate starvation and increases again when the seedlings are transferred back to low nitrate conditions (Figure 1E). This time, we also found an impact on *NRT2.1* inter-individual transcriptional variability, that mimics what is observed for *NRT2.1* expression (Figure 1F): the variability is strongly reduced when plants face complete nitrate starvation and increases again after plants are back to low nitrate conditions.

All in all, these results show that *NRT2.1* is a highly variable gene in several environmental conditions, except in the case of a complete nitrate starvation where the loss of the variability for *NRT2.1* is likely caused by the loss of its expression.

### *NRT2.1* inter-individual transcriptional variability is associated with phenotypic variability

To define how meaningful *NRT2.1* inter-individual variability might be, we compared the differences in expression between plants for *NRT2.1* with the transcriptional response of that gene to the environment (Figure 2A). We found for the *pNRT2.1:LUC* reporter line, that the difference in the average signal for seedlings grown in low (1mM KNO_3_) or high (10mM KNO_3_) nitrate conditions (1.7x between the medians) is in the same order as differences between seedlings in a given condition (1.4 to 1.9x between the 1^st^ and the 3^rd^ quartile, Figure 2A). The differences in expression between these two environmental conditions are known to have a physiological consequence, NRT2.1 being the main nitrate transporter in low nitrate conditions while other transporters, like NRT1.1, are involved for plants in high nitrate conditions (Wang et al., 2012; O’Brien et al., 2016). In this context, our result suggests that the extent of *NRT2.1* transcriptional variability between plants might be relevant since the level of differences between plants in one environment is at least as much as the transcriptional response of that gene to changes in nitrate concentrations.

**Fig 2.**
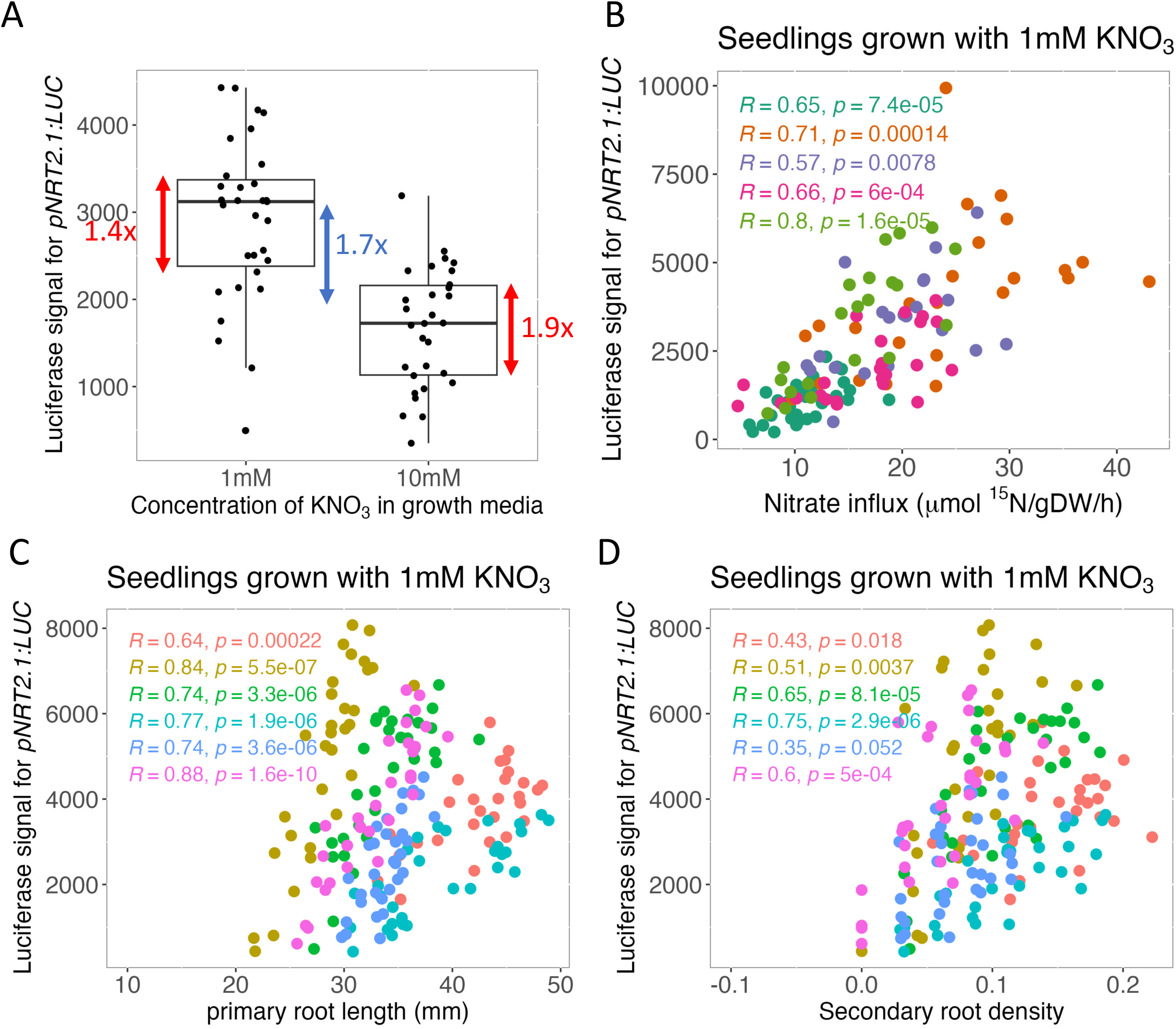
Phenotypic consequences of plant-to-plant *NRT2.1* variability. (A) Comparison for the *pNRT2.1:LUC* line of the differences in signal between seedlings in one condition (1.4x to 1.9x in red) and between the mean signal of seedlings grown in media with low (1mM) of high (10mM) nitrate concentration (1.7x in blue). Each point corresponds to the signal in a single seedling. (B-D) Comparison for plants grown in media with low (1mM) nitrate concentration of the *pNRT2.1:LUC* signal and (B) the nitrate influx of HATS, (C) the primary root length or (D) the lateral root density. Each point corresponds to a single seedling and each colour to a replicate. The result of a spearman correlation test for each replicate is included.

To test if *NRT2.1* variability between plants could indeed have phenotypic consequences we co-analysed in the same seedlings the level of *NRT2.1* promoter activity using the *pNRT2.1:LUC* reporter line and several phenotypes known to be affected in the *nrt2.1* mutant: the activity of the high affinity nitrate transport system (HATs), the primary root length as well as the lateral root density (Cerezo et al., 2001; Wang et al., 2023; Little et al., 2005; Remans et al., 2006). First, we measured the luciferase signal for individual seedlings and then performed a nitrate influx experiment to measure the activity of HATs in the same seedling. Since *NRT2.1* explains most of the transport observed for HATs when plants are in low nitrate conditions (Cerezo et al., 2001; Filleur et al., 2001), our measurement can be considered as a proxy for *NRT2.1* protein activity. We observed a strong positive correlation between the luciferase signal for *pNRT2.1:LUC* and the nitrate influx for HATs in all 7 replicates when seedlings are grown on low nitrate (1mM KNO_3_) conditions (Figure 2B, Figure S2B, spearman p-value < 0.05). However, this correlation is lost in all 4 replicates when seedlings are grown on high nitrate (10mM KNO_3_) conditions (Figure S2A-B). This result suggests a phenotypic consequence of *NRT2.1* transcriptional variability, seen via the correlation with the variability of NRT2.1 transporter function, only when the plants are grown in conditions where NRT2.1 is the main transporter.

To explore the possible impact of *NRT2.1* transcriptional variability on macroscopic phenotypes representative of plant growth and development, we did the same analysis for the primary root length and the lateral root density. We found a strong positive correlation in all 6 replicates between *NRT2.1* promoter activity reported by the luminescence measured along the primary root, and the primary root length measured on single seedlings for plants grown with low (1mM KNO_3_) (Figure 2C) as well as high (10mM KNO_3_) concentrations of nitrate (Figure S2C). We also observed a positive correlation between *NRT2.1* promoter activity in the primary root and the lateral root density measured on single seedlings, but this mainly for plants grown with a low (1mM KNO_3_) nitrate concentration (Figure 2D). For plants grown with a high (10mM KNO_3_) nitrate concentration, the correlation with the lateral root density was not as strong and only significant for 4 out of 6 replicates (Figure S2D). As a control, we did not observe a correlation between the *p35S:LUC* signal and the primary root length in both low and high nitrate concentrations, except in one of the three replicates (Figure S2E-F).

These results suggest a possible phenotypic consequence of *NRT2.1* inter-individual transcriptional variability. To verify if this could be the case, we measured the impact of the *nrt2.1* mutation on the primary root length and the lateral root density in our conditions. We found that *nrt2.1* mutant has a shorter root length and a lower lateral root density than the WT when they are grown on low nitrate conditions (1mM KNO_3_) (Figure 3A-B). This is in agreement with the positive correlation we measured between these two root phenotypes and the luciferase signal for *pNRT2.1:LUC*, showing that plants with a high promoter activity for *NRT2.1* have a longer primary root and a higher density of lateral roots. However, no reproducible difference between the two genotypes was observed if the plants are grown on high nitrate conditions (10mM KNO_3_) (Figure 3B).

**Fig 3.**
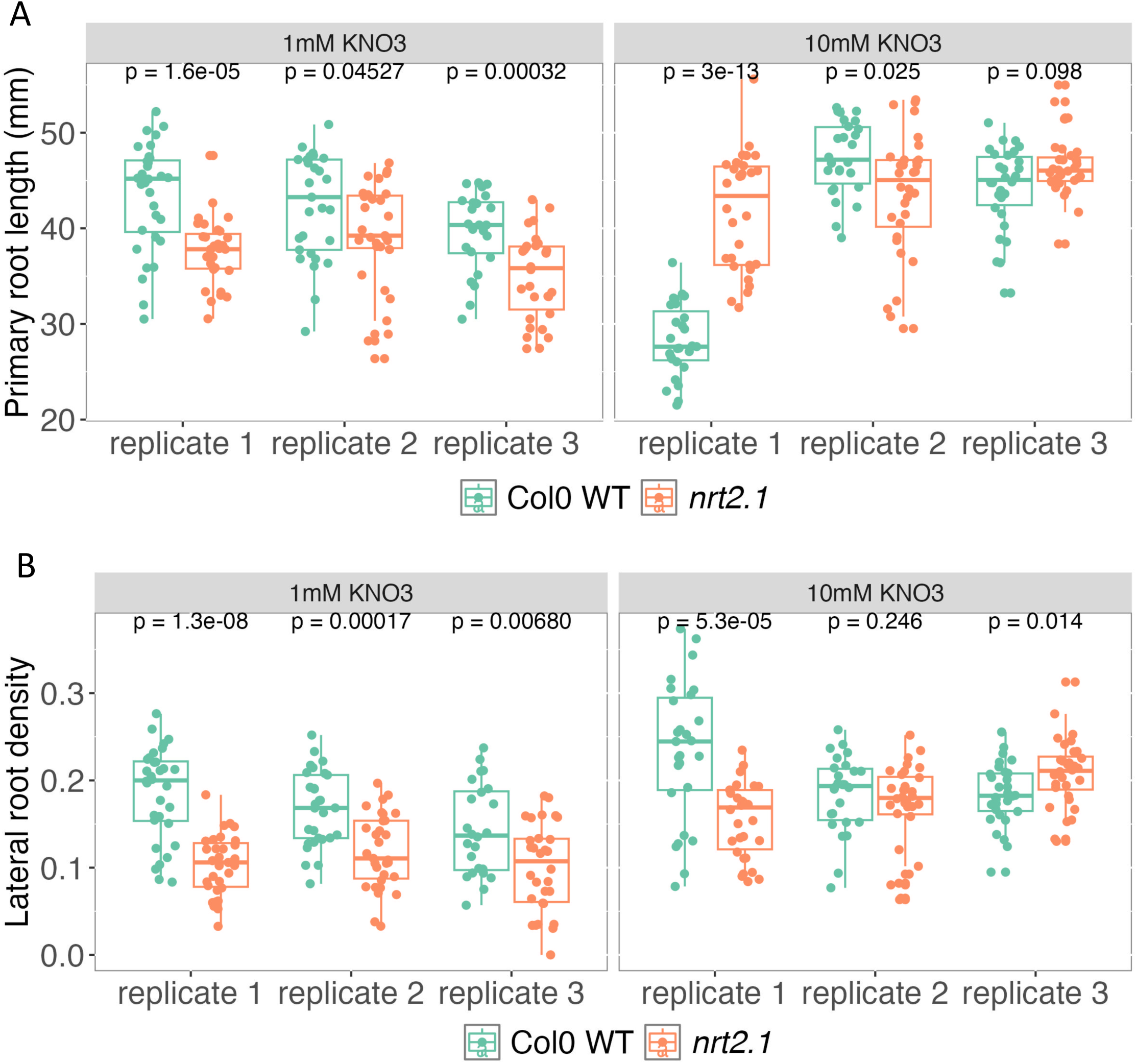
Impact of *nrt2.1* mutation on root phenotypes. (A-B) Measurement of (A) the primary root length and (B) lateral root density (number of lateral roots/ primary root length) for Col0 WT and the *nrt2.1* mutant for plants grown with either 1mM of KNO_3_ (left) or 10mM of KNO_3_ (right) in their growth media. A total of 3 biological replicates are shown. The results of a Wilcoxon test are included.

All in all, our results imply that inter-individual transcriptional variability for *NRT2.1* is likely the cause for phenotypic variability, for the primary root length and lateral root density, as well as NRT2.1 promoter and protein activities, measured through the nitrate influx for HATs, when plants are grown in low nitrate conditions. For these reasons, all the rest of the study will be performed on plants grown with low nitrate concentrations (1mM KNO_3_).

### *NRT2.1* expression level in a plant is maintained over days but not over generations

To understand how the inter-individual transcriptional variability for *NRT2.1* is established, we first need to better characterize it. We thus decided to define if *NRT2.1* expression level in a plant is conserved over days or if there is a daily reset. Indeed it is known that *NRT2.1* expression is dependent on light and is thus only expressed during the day but not during the night (Lejay et al., 1999). For this, we analysed the luciferase signal at 10, 14, 17 and 21 days for *pNRT2.1:LUC* seedlings growing on low nitrate conditions. We observed that, in general, seedlings keep their relative *NRT2.1* expression level throughout the experiment (Figure S4A). Moreover, there was a significant positive correlation between the signal of seedlings at the first and last day of the experiment (Figure S4B). This shows the absence of a daily reset for *NRT2.1* expression level in seedlings. To ensure these differences in expression are not caused by the microenvironment around the seedlings, we transferred 6-day-old *pNRT2.1:LUC* seedlings to a new plate containing the same condition (low nitrate) and measured their luciferase signal just before the transfer and for 4 days after transfer. We found that, despite the transfer of seedlings to a new plate, most of them kept their relative expression levels during the experiment (Figure 4A-B). Seedlings with a high expression before transfer still have a high expression when in the new plate and vice-versa. Moreover, there is a positive significant correlation of the luciferase signal in seedlings before and 4 days after transfer (Figure S4C). In both experiments, most seedlings kept their relative expression levels as compared to other plants, and only a minority of individuals changed drastically their expression level upon transfer. These results indicate that the differences in *NRT2.1* expression level between seedlings are not caused by the microenvironment but by a state of the seedling that is maintained over several days.

**Fig 4.**
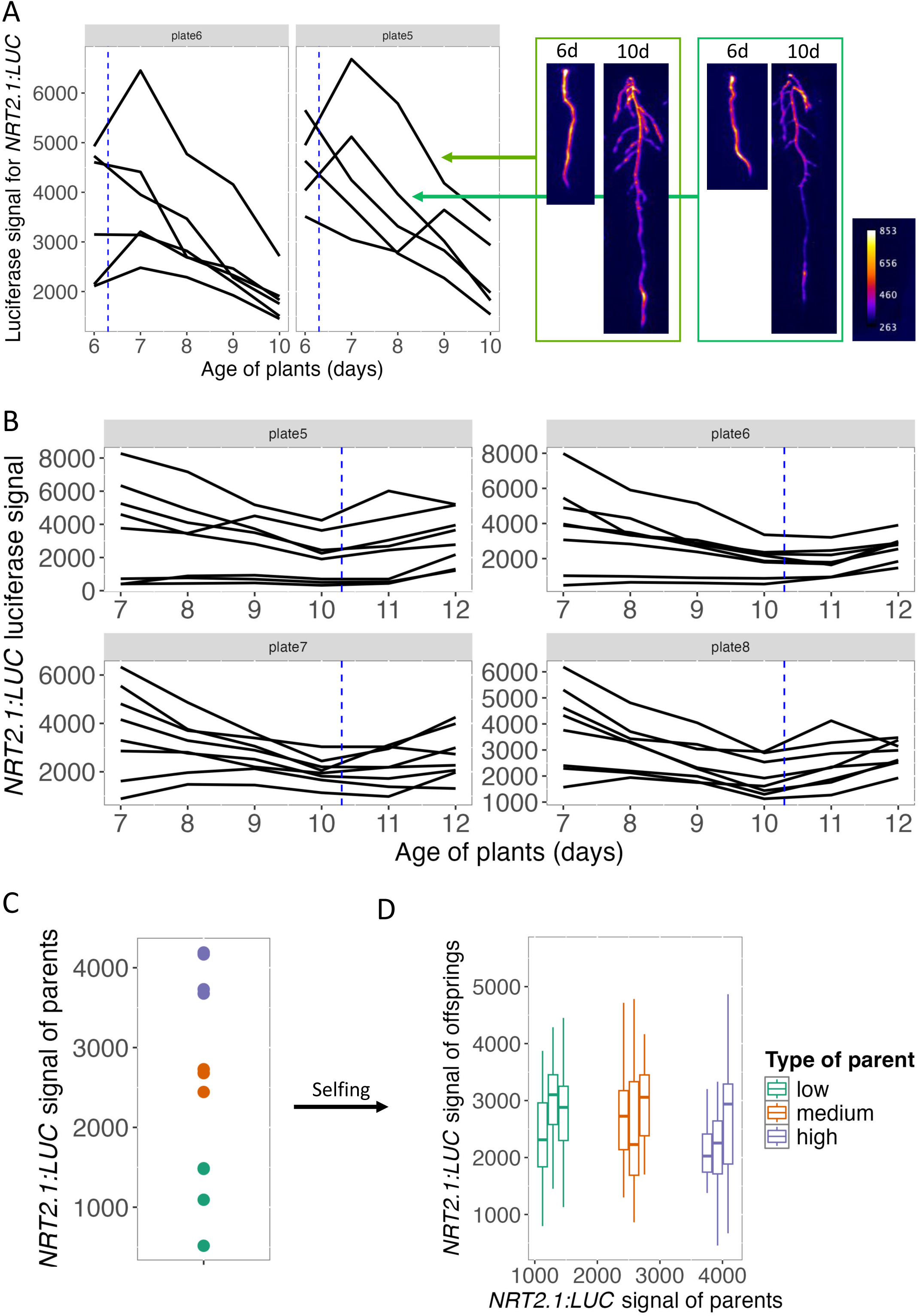
*NRT2.1* expression levels are stable over time but not transmitted at the next generation. (A) *pNRT2.1:LUC* signal over 5 days, after a transfer to a new plate, for two plates. Each line corresponds to a given seedling. The vertical blue dashed line indicates the transfer of seedlings to a new plate. (B) *pNRT2.1:LUC* signal over 6 days, with a transfer to a new plate in the middle of the experiment, for four plates. Each line corresponds to a given seedling. The vertical blue dashed line indicates the transfer of seedlings to a new plate. (C) *pNRT2.1:LUC* signal for 19 seedlings. The individuals selected to analyse the signal of their offsprings are shown in colour depending on their category: low expression in green, medium expression in orange and high expression in purple. (D) Distribution of the *pNRT2.1:LUC* signal for populations deriving from self-pollination of the parents selected in (B). Each boxplot corresponds to a population of descendants (at least 24 seedlings), with the colour depending on the category of the parent: low expression in green, medium expression in orange and high expression in purple.

To go further, we wanted to test if the relative expression levels of seedlings are maintained when facing environmental fluctuation. For this, we analysed the starvation experiment already used above and compared the *pNRT2.1:LUC* luciferase signal in individual seedlings before and after a complete starvation. We observed a significant positive correlation for the signal in individual seedlings prior and after starvation (Figure S4D). This result, as well as the maintenance of expression levels over days, indicate the relative expression for *NRT2.1* in each seedling is maintained even after a reset of its expression, which are caused by these two conditions (dark in one case, nitrate starvation in the other). All in all, it thus seems that relative expression levels between seedlings is independent of the transcriptional regulation of *NRT2.1* in response to the environment.

Since we have seen that *NRT2.1* expression level in seedlings are stable over several days, we then wanted to test if they could be transmitted to the next generation. For this, we selected several *pNRT2.1:LUC* seedlings that had a high, medium or low *NRT2.1* expression level (Figure 4C, Figure S4E), transferred them to soil and let them produce seeds. We then analysed the luciferase signal in the descendants to define if their expression level is influenced by the one in their parents. If there is a transmission of *NRT2.1* expression level, we expect descendants of a parent with high *NRT2.1* expression level to also display a high expression and the ones coming from a parent with low *NRT2.1* expression level to have a low expression. We found a similar *NRT2.1* expression level in all populations of descendants irrespective of the expression level in their parents, and this for two independent experiments (Figure 4D, Figure S4F). This result indicates that *NRT2.1* expression level in each seedling is not transmitted to the next generation. It suggests a reset in the regulation of *NRT2.1* and that differences in expression between seedlings are re-established at each generation.

### Differences in expression between seedlings are established in young seedlings

The next step is to define when the differences in expression between plants are established. To do this, we followed *NRT2.1* promoter activity every 4 hours for 10 days from the exit of stratification using the *pNRT2.1:LUC* line. The *p35S:LUC* line was also analysed in parallel as a control. This was done using an automated plant growth chamber with controlled light phases for growth and dark phases with luminescence imaging, in which up to 11 plates can be analysed in the same experiment (Lumalum, Hanchi et al., 2018). Luminescence signals were again measured along the primary roots. For *pNRT2.1:LUC,* we observed a transient peak of signal at germination, followed by a gradual increase of signal over days (Figure 5A, Figure S5). At around seven to eight days, we observed an increase in the difference for *pNRT2.1:LUC* signal in the different seedlings. In some plates it was very striking (see plates 0 and 4), with the signal that continued to increase in some seedlings, while it stagnated or even went down in others. From that moment, seedlings will usually maintain a high or a low *pNRT2.1:LUC* signal until the end of the experiment (Figure 5A-B, Figure S5). This is different from what is observed for the *p35S:LUC* line where the signal increases gradually in a similar way in all seedlings (Figure 5A, Figure S5) These results indicate that differences in expression between seedlings for *NRT2.1* are established at around 7 days and are then maintained over time.

**Fig 5.**
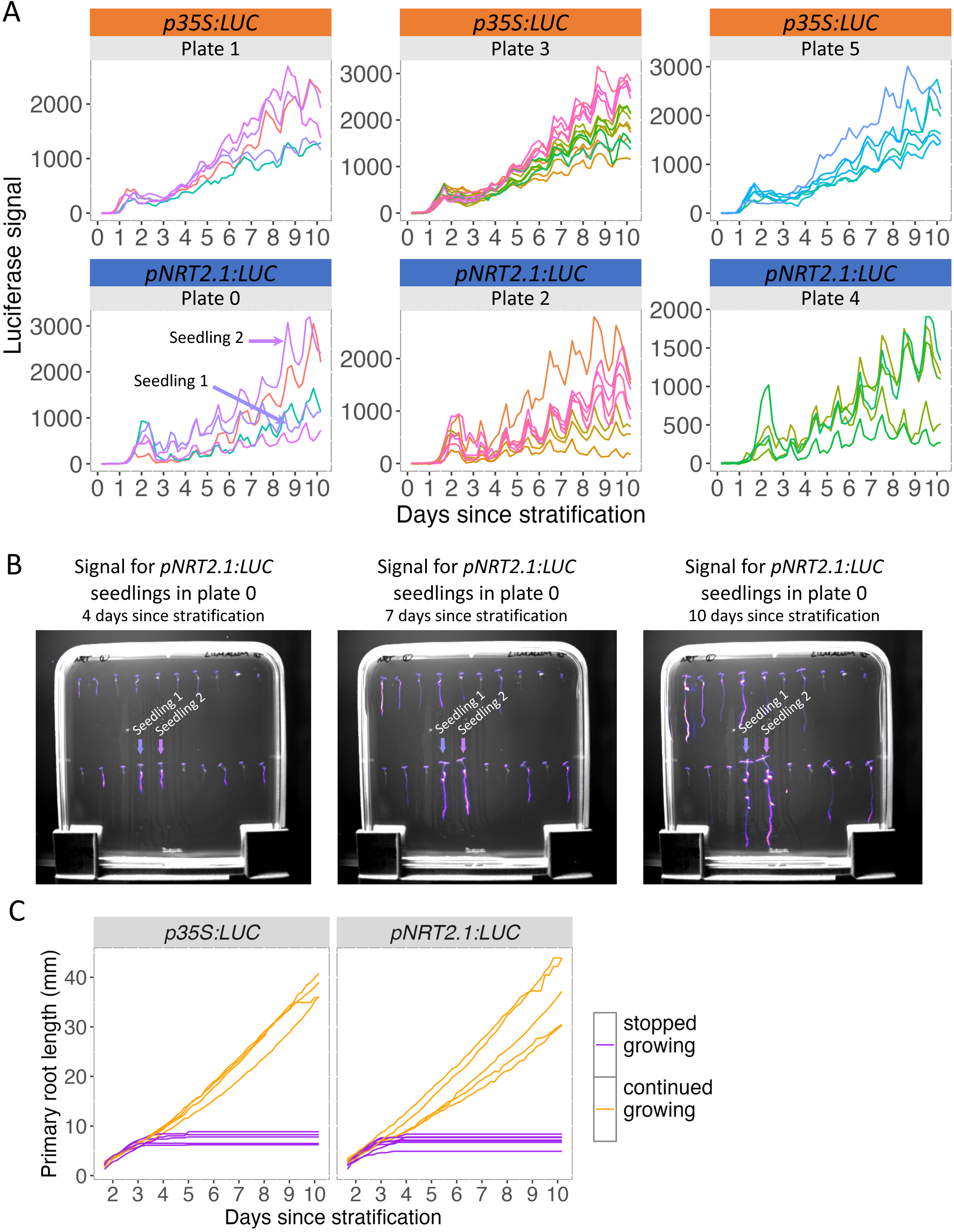
Differences in expression between seedlings for *NRT2.1* are established in young seedlings. (A) *p35S:LUC* (top) and *pNRT2.1:luc* (bottom) *s*ignal over time for 3 plates of each genotype. Five to twelve seedlings were measured in each plate. Only seedlings that have lateral root at the end of the experiment were measured. Each line represents the signal in a given seedling, measured every 4 hours for 10 days after stratification. (B) Images of signal for pNRT2.1:LUC seedlings in the plate 0 at four, seven and ten days after stratification. The colour scale of the signal is the same in all images. Seedling with low signal (seedling 1) or high signal (seedling 2) at the end of the experiment are indicated. (C) Primary root length during the time course, measured every 4 hours from one day to ten days after stratification. Seedlings with roots that stop growing and fail to do the heterotroph to autotroph transition are shown in purple while seedlings with roots that continue to grow throughout the entire experiment are shown in yellow.

To test if differences in *NRT21* expression are established before or after the heterotrophic-to-autotrophic transition in young seedlings, we analyzed the primary root growth of seedlings from germination. We observed in some seedlings a primary root growth arrest, very likely due to a failure in the heterotrophic-to-autotrophic transition (Kircher and Schopfer, 2012), happening at around 3 days (Figure 5C).

All in all, our results show that differences in NRT2.1 expression between plants are established in young seedlings, after the heterotrophic-to-autotrophic transition, and then maintained over days.

### New factors involved in regulating *NRT2.1* expression level in seedlings

In order to go further and understand better what might be the cause of the inter-individual transcriptional variability for *NRT2.1* we carried out a transcriptomic study. For this, we performed RNA-seq on 24 individual seedlings that were grown on a media with low nitrate concentration (1mM KNO_3_). By comparing the transcriptome of the seedlings, we can then measure the level of inter-individual transcriptional variability for all genes (Cortijo et al., 2019). Focussing on the genes involved in nitrate nutrition, we see that *NRT2.1* is highly variable, as well as *PP2C* (A type 2C protein phosphatase), involved in NRT2.1 protein regulation, and some of its negative transcriptional regulators: *BT1*, *BT2*, *HHO3*, *HHO2* and *LBD37* (Ohkubo et al., 2021; Araus et al., 2016; Kiba et al., 2018; Rubin et al., 2009). On the other hand, none of the positive transcriptional regulators of *NRT2.1* and the genes involved in nitrate assimilation are highly variable (Figure 6A). This result suggests that the high inter-individual variability for some of the genes involved in the negative transcriptional regulation of *NRT2.1* might be the cause of the variability for that gene.

**Fig 6.**
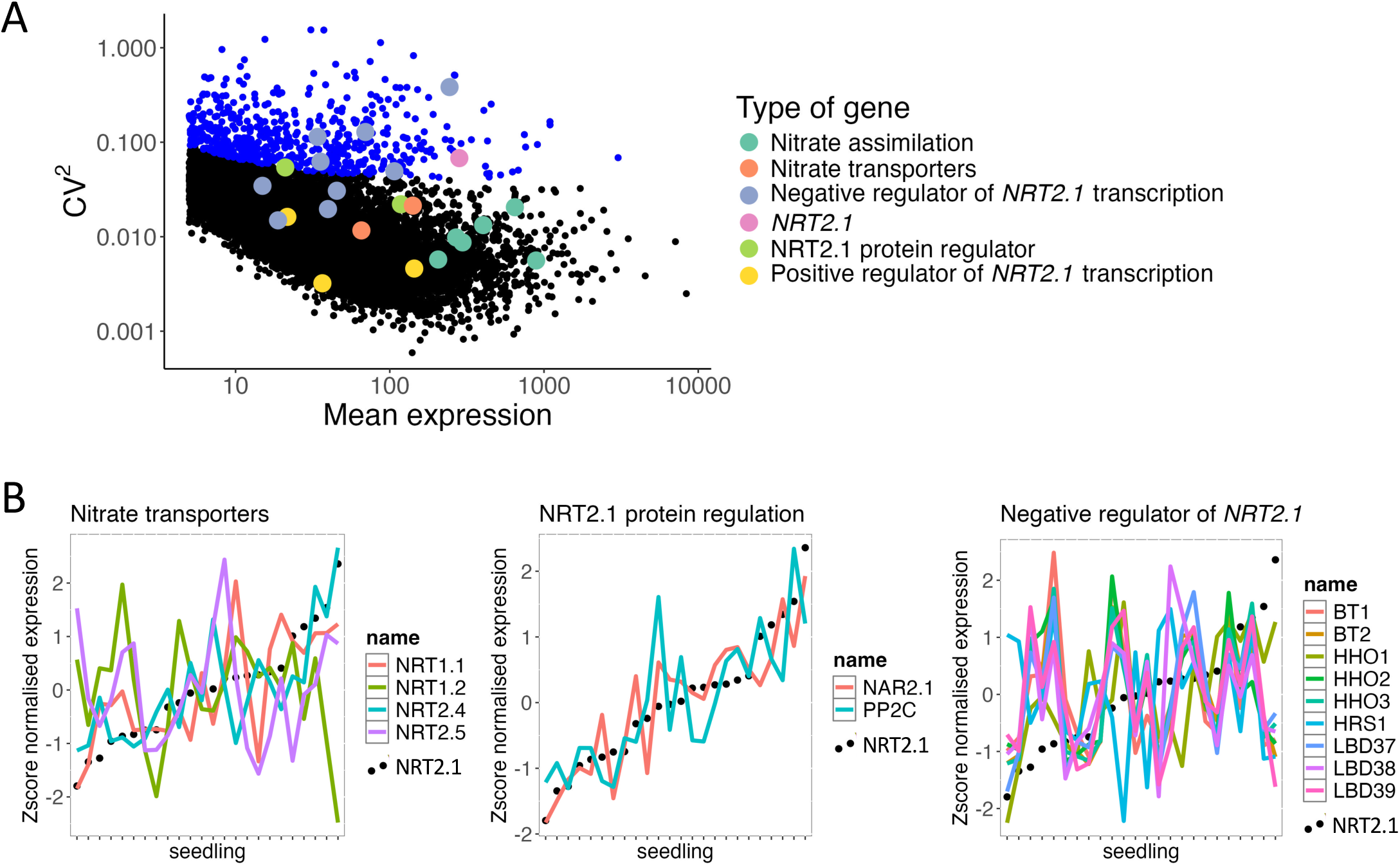
*NRT2.1* inter-individual transcriptional variability is not associated with its transcriptional regulators. (A) Representation for the entire transcriptome of the CV against the mean expression, measured on 24 seedlings. Each point represents one gene, and the points in blue indicate statistically highly variable genes. The genes involved in different aspects of nitrate nutrition are shown in different colours. (B) Expression level in the different seedlings of *NRT2.1* (black dots) and nitrate transporters (left), genes involved in the post-translational regulation of *NRT2.1* (middle) and negative regulators of *NRT2.1* expression (right).

To test this, we compared the expression level of *NRT2.1* and its regulators in the different seedlings to define if they are correlated (Figure 6B, Figure S6A and table S1). We saw that the two genes involved in the regulation of NRT2.1 protein activity, *NAR2.1* and *PP2C* (A type 2C protein phosphatase) (Laugier et al., 2012; Ohkubo et al., 2021; Orsel et al., 2007), are expressed in a very similar way as *NRT2.1* in the different seedlings (p-value <0.05 and R^2^>0.7, spearman correlation test). This could explain the positive correlation we observed between NRT2.1 promoter and protein activities in single seedlings (Figure 2B). For other categories of genes (nitrate transporters, nitrate assimilation, positive or negative transcriptional regulators of *NRT2.1*) we did not observe such a positive correlation in expression in the seedlings between *NRT2.1* and all genes of each category. However, some genes of these categories had a very similar expression as *NRT2.1* in the seedlings: the transporters *NRT2.4* and *NRT1.1* as well as the nitrite reductase (*NIR1)* (p-value <0.05 and R>^2^0.7, spearman correlation test). In particular, we could not see a strong correlation between *NRT2.1* and any of the genes involved in its transcriptional regulation (Figure 6B, Figure S6A and table S1). Since it is known the transcription factor NLP7 is not regulated transcriptionally but at the post-translational level (Marchive et al., 2013), we compared the expression in seedlings between *NRT2.1* and the targets of NLP7 and did not found any correlation (Figure S6B). These results suggest the inter-individual transcriptional variability for *NRT2.1* might not be caused by changes in expression between seedlings for the genes known to regulate its expression.

To find other pathways that might be at the origin of *NRT2.1* inter-individual transcriptional variability, we performed a hierarchical clustering of all genes that vary sufficiently in expression between seedlings (CV>1, measured using the expression level in all seedlings). We identified 6 clusters of genes that have different types of expression profiles in the seedlings. Among them, the cluster 1 which is composed of 391 genes contains *NRT2.1* (Figure 7A). Globally, the genes of this cluster 1 have an expression in the different seedlings relatively similar to *NRT2.1* (Figure S6C). We thus performed a gene ontology enrichment for the genes in this cluster, and surprisingly, we didn’t see any enrichment for gene ontologies associated with nitrate or N nutrition, but mainly with carbon, light and photosynthesis as well as development (Figure 7B). Since it is know that nitrate and carbon metabolisms are coordinated in plants (Martin et al., 2002), we wanted to explore their relation in our data. For this, we analysed the correlation between *NRT2.1* expression in single seedlings and genes that have been shown to be involved in different energy associated pathways as previously described (Avin-Wittenberg et al., 2012). Among them are genes involved in photosynthesis, tetrapyrrole biosynthesis, glycolysis, TCA cycle, Asp-family pathway and ATP biosynthesis. We found the highest proportion of genes for which the expression in single seedlings is statistically correlated with *NRT2.1* expression for the genes involved in photosynthesis (48%, Figure 7C). This is higher than for genes involved in nitrate nutrition (32%).

**Fig 7.**
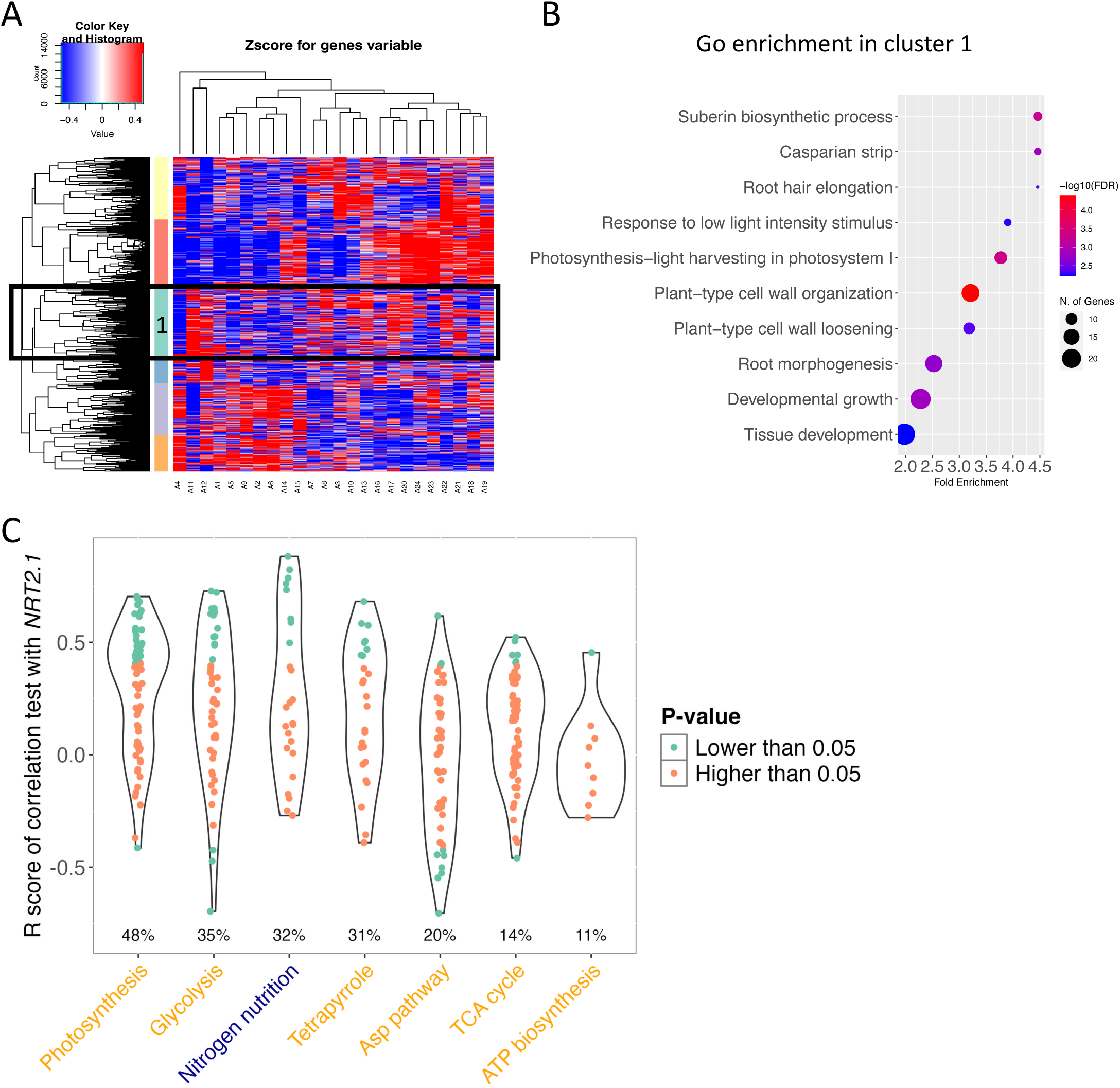
*NRT2.1* inter-individual transcriptional variability seems to be associated with primary metabolism. (A) Hierarchical clustering in the different seedlings of genes with a corrected CV>1. The Z-score is used to correct for expression and variance levels. *NRT2.1* is in the cluster 1 (light blue cluster). (B) Gene ontology enrichment analysis for the cluster 1 which contains *NRT2.1*. The top 10 gene ontologies enriched are represented. (C) Rscore of the correlation with *NRT2.1* expression in single seedlings for genes involved in different pathways: photosynthesis, glycolysis, tetrapyrrole, Asp pathway, TCA cycle and ATP biosynthesis. Genes involved in nitrogen nutrition are also included for comparison. The different pathways are ordered based on the proportion of the genes for this pathway with a p-value less than 0.05 (highest on the left, lowest on the right).

Altogether, these results indicate that genes co-expressed with *NRT2.1* in the different seedlings are not associated with nitrate nutrition but with carbon metabolism and development. It suggests that new factors and pathways, yet to be identified, coordinate *NRT2.1* expression with genes involves in carbon metabolism, and their inter-individual variability.

## Discussion

In this study we found that *NRT2.1* gene expression is highly variable between genetically identical plants grown in the same environmental condition (Figure 1). The only condition where we observed a loss of *NRT2.1* inter-individual transcriptional variability was when plants were subjected to a complete nitrate starvation, in which case the expression of *NRT2.1* was also lost (Figure 1). Interestingly, the differences in expression between plants for *NRT2.1* were correlated with several phenotypes: the influx of nitrate by high affinity transporter system as well as the primary root length and lateral root density (Figure 2). We also found that while the expression level of *NRT2.1* in a seedling is usually maintained for at least several days, it is not transmitted to the next generation (Figure 4). This led us to wonder what is the source of this high inter-individual variability. To explore it, we decided to define when differences in expression between plants are put in place. We found that *NRT2.1* expression diverges between plants in young seedlings after the heterotrophic-to-autotrophic transition (Figure 5). To go further we also analysed genes that are co-expressed with *NRT2.1* at the genome-wide scale. We found that known transcriptional regulators of *NRT2.1* are not co-expressed with it, and observed on the contrary an enrichment for functions related to root growth as well as photosynthesis among genes co-expressed with *NRT2.1* (Figures 6-7).

When exploring the possible phenotypic impact of inter-individual transcriptional variability, we found that differences in *NRT2.1* promoter activity between seedlings are correlated with the high affinity root nitrate transport (Figure 2). This is in agreement with previous studies done on bulks of plants showing a strong reduction of the nitrate influx by high affinity root nitrate transport in the *nrt2.1* mutant (Remans et al., 2006; Little et al., 2005). However, it is also known that *NRT2.1* expression levels do not systematically correlate with the amount of NRT2.1 protein and that it might be subjected to post-translational regulation (Wirth et al., 2007). For example, PP2C (AT4G32950), a type 2C protein phosphatase, was shown to dephosphorylate NRT2.1 protein and activate it (Ohkubo et al., 2021). Moreover, NRT2.1 function is also regulated by its interaction with NAR2.1 (AT5G50200), a small transmembrane protein (Orsel et al., 2007; Laugier et al., 2012). Interestingly, we found that the expression of the genes for these two proteins is positively correlated with *NRT2.1* expression in single seedlings (Figure 6). It suggests these two regulators and *NRT2.1* itself are in the same regulatory pathway when it comes to the control of their expression, and that the level of activation of this pathway is different from one plant to another. In light of these results, it is thus very likely that differences in expression between seedlings for *NRT2.1* as well as its two regulators *PP2C* and *NAR2.1* are the cause of the phenotypic variability for high affinity root nitrate transport for plants grown in low nitrate conditions.

We also found a strong positive correlation between *NRT2.1* promoter activity and two root phenotypes: primary root length and the density in lateral roots (Figure 2). In addition, we showed that the *nrt2.1* KO mutant has a shorter primary root and a lower lateral root density than the WT when analysed in the low nitrate conditions (1mM KNO_3_) (Figure 3). This is also supported by the fact that GO involved in root growth and development are enriched among the genes co-expressed with *NRT2.1* in our single seedling RNA-seq (Figure 7). When it comes to primary root length, our results are different from what was previously published, where a longer primary root length was observed in the *nrt2.1* mutant compared to the WT when grown on low nitrate condition (Wang et al., 2023). However, the growth media contained 1% sucrose in this previous study, while we do not provide any. Since it is known that the carbon/nitrogen balance has an important role in regulating the growth of plants (Martin et al., 2002), the presence or absence of sucrose in the media in the 2 studies could explain the opposite results. On the other hand, our results for the density of secondary roots are supported by a previous study analysing the *nrt2.1* mutant in low nitrate condition in absence of sucrose (Remans et al., 2006). Again, a different result was observed in another study done with sucrose in the media (Little et al., 2005). Altogether, our results indicate that plant-to-plant variability for primary root length and lateral root density are likely caused by the differences in *NRT2.1* expression between plants. A possibility would be that *NRT2.1* expression level in a given plant affects its global growth rate. Supporting this hypothesis, several studies show that *nrt2.1* mutant is smaller than WT, even if they mainly focus on the aerial part or the total weight of the plant and not the root area (Lezhneva et al., 2014; Kiba et al., 2012). However, we cannot completely rule out that differences in growth rates between plants could change their internal nitrogen status (concentration of nitrogen metabolites), which in turn has been shown to regulate the expression level of *NRT2.1* (Lejay et al., 1999; Zhuo et al., 1999). In order to discriminate between these two possibilities, we would need to reduce or increase *NRT2.1* inter-individual transcriptional variability, without affecting its average expression level, and see if it also affects root growth.

We found that *NRT2.1* has a high inter-individual transcriptional variability in several environmental conditions tested. Moreover, we found that *NRT2.1* expression level and variability could be uncoupled. For example, *NRT2.1* inter-individual transcriptional variability is not significantly different when plants are grown in low or high nitrate even if the overall level of expression for that gene is different (Figures 1-S1). This is in agreement with a previous study where *NRT2.1* was detected as a highly variable gene for plants growing on MS/2, where the expression for this gene is very low (Cortijo et al., 2019). The only case where we observed a loss of variability is during a complete nitrate starvation (Figure 1), a quite significant stress for the plants. Interestingly, it was previously observed in yeast that genes with a high variability in normal conditions showed a reduction of their variability in response to stress (Gasch et al., 2017). This study, done at the transcriptome level, also showed that genes that are normally lowly variable also displayed an increase in variability during stress. Another study in *Escherichia coli* of the response of gene expression level and noise in different environmental conditions (including some stresses) shows a variety of relations between noise and expression level (Urchueguía et al., 2021). They found positive as well as negative relations between the changes in noise and expression levels in different environments for some genes while for other genes the level of noise didn’t change whatever the conditions even if the expression level changed. It shows no global link between changes in expression and noise in response to the environment, but more a gene by gene relation. This is in agreement with what was observed in a study of inter-individual transcriptional variability and expression level in plants throughout a day/night cycle (Cortijo et al., 2019). All possibilities were observed when it comes to the relation between transcriptional variability and expression level (positive or negative relation as well as none). The only study of transcriptional noise during stress in plants was done in response to heat stress for a couple of heat responding genes but was focussed on cell-to-cell noise and did not analyse inter-individual transcriptional variability. They showed a high cell-to-cell noise for these genes in response to heat stress (Alamos et al., 2021). However, they were not expressed prior to stress, making it difficult to compare their level of noise before and after the stress. There are currently no studies of inter-individual transcriptional variability in response to stress in plants. To see if our result could be generalised, a genome-wide study of inter-individual transcriptional variability and expression level for plants subjected to several stresses will be needed. Only this type of experiment will allow us to better understand the relation between the response to stresses for gene expression and inter-individual transcriptional variability.

We observed that differences between plants for *NRT2.1* expression level are established in young seedlings after the heterotrophic-to-autotrophic transition (Figure 5). At this point in life, plants transition from using the reserves accumulated during seed development (triacylglycerols and seed storage proteins in *Arabidopsis thaliana*) to generate their own energy via photosynthesis (Tan-Wilson and Wilson, 2012; Graham, 2008). This is a period characterized by massive changes in the regulation of the transcriptome, the epigenome and the condensation of the chromatin (Silva et al., 2016; Simon and Probst, 2024; Samo et al., 2024). We could suppose that during this transition, the regulation of genes involved in nitrate nutrition is coordinated with carbon assimilation. This is in agreement with the fact that, in our transcriptomic analysis, we found that genes involved in photosynthesis, in nitrate assimilation and NRT2.1 protein regulation, but not its transcriptional regulation, are strongly co-expressed with *NRT2.1* in seedlings (Figure 7). Together, it suggests a coordinated regulation of sets of genes allowing a good carbon/nitrogen balance in plants when photosynthetic activity is established. After that period in young seedlings where differences in expression for *NRT2.1*, and possibly other genes involved in carbon metabolism, are established, they are maintained over time.

We will need to study the dynamics of expression at a single plant resolution in young seedlings for other genes, especially ones involved in very different functions, to define if *NRT2.1* is a special case or if differences in expression between plants usually happen shortly after the heterotrophic-to-autotrophic transition. For this, auto-bioluminescent reporters with multiple colours, that have been developed recently in plants (Kusuma et al., 2024; Shakhova et al., 2024), will allow us to follow the promoter activity of multiple genes in the same plants. Moreover, to understand how differences in expression are established between young seedlings, we would need to explore more in detail the transcriptional and chromatin regulations happening. For now, the only data available were generated using pools of seedlings are available (Silva et al., 2016; Simon and Probst, 2024; Samo et al., 2024) and while they improve our knowledge of the transcriptomic and epigenomic dynamics happening during the heterotrophic-to-autotrophic transition, they mask any plant-to-plant differences. Analysing the transcriptomes and epigenomes of single seedlings before and after the heterotrophic-to-autotrophic transition will allow us to define if there is any coordinated regulation of genes and the possible role of chromatin in the establishment of inter-individual transcriptional variability.

## Material and methods

### Plant material and growth conditions

All lines used in this study are in the Col-0 background. The *pNRT2.1:LUC* line and *p35S:LUC* lines, already published (Girin et al., 2010; Dorbe et al., 1998), contain 1200bp of *NRT2.1* promoter and the 35S promoter fused with the luciferase gene, respectively. The *nrt2.1* KO mutant is the SALK_035429 T-DNA insertion line.

For all experiments, plants were grown on petri dishes after seed sterilisation and stratification, on a solid media containing 1mM KH_2_PO_4_, 1mM MgSO_4_, 0.25mM K_2_SO_4_, 0.25mM CaCl_2_, 0.1mM Na-Fe-EDTA, 2.5mM MES and microelements (50μM KCl, 30μM H_3_BO_3_, 5μM MnSO_4_, 1μM ZnSO_4_, 1μM CuSO_4_ and 0.1μM (NH_4_)_6_Mo_7_O_24_). Only the concentration of KNO_3_ differs depending on the experiment: 1mM KNO_3_ (low nitrate), 10mM KNO_3_ (high nitrate) or 0mM KNO_3_ (no nitrate). Except for the time-course experiments with the Lumalum, a total of 4 plates per genotype and condition were prepared, with 30 seeds per plate (disposed over 2 rows). After 6 days of growth on long days at 22 degrees Celsius and a light intensity of 150 μmol m^−2^ s^−1^, seedlings that managed to do the heterotrophic-to-autotrophic transition and continued to grow were transferred to a new plate. The KNO_3_ concentration of the media in the first and second plate are specified in case of a specific treatment, and otherwise stay the same if not specified. After transfer, seedlings were put back in the growth chamber. Luciferin at a final concentration of 0.25 mM was also included in the growth media for all luciferase experiments.

### Luciferase imaging

Except when specified, luciferase imaging was done on 9-day-old seedlings. Most of the snapshot experiments where done using a cooled Hamamatsu C4880-30-24W CCD camera with a 2 minutes acquisition time following a 5 minutes period of dark in order to remove any auto-luminescence caused by photosynthetic activity. Because this camera broke close to the end of the project, imaging for the figure S2B was done with another camera: a cooled Andor iXon ultra 897 EMCCD Back-illuminated with a 20 seconds of acquisition time following a 5 minutes period of dark.

For time-course luciferase imaging over several days from germination, a dedicated plant growth chamber with controlled light and dark phases as well as automated luminescence imaging (Lumalum) was used (Hanchi et al., 2018). Temperature inside the imaging growth cabinet was maintained constantly at about 25°C +/− 0.5°C to limit the impact of temperature on luciferase enzymatic activity. Light intensity of the LED module was adjusted to about XµE, with respective LED intensities of 2% for 420nm, 20% for 450n, respectively. A total of 11 plates were analysed at the same time. Throughout the time-course, luciferase imaging was performed every four hours on all plates, with a 60 seconds (bin2) acquisition time. Reference bright field images were taken with a 200 msec exposition (bin2).

The raw integrated density (sum of the signal for all pixels) for luciferase was measured along the primary root (not along the lateral roots) of each seedling using Fiji (Schindelin et al., 2012) and then corrected for the background signal of the same plate as well as the length of that primary root.

### RNA extraction and RT-qPCR

RNA was extracted from 11-day-old individual seedlings using the MagMAX™-96 Total RNA Isolation Kit (Ambion) following manufacturer’s instructions except for two changes: 60µl of Lysis/Binding solution were used instead of 100µl, and the elution was performed with 30µl of Elution Buffer instead of 50µl. One microgram of RNA was used for reverse transcription, using the M-MLV Reverse Transcriptase (Invitrogen) following manufacturer’s instructions. qPCRs were performed in 384 well plates with 10µl reactions using a LightCycler 480 (Roche). Primers used in this study are: CAACTACAACCTCACGCAGC and ACCCTTCTTATTCCTCCGGC for the lowly variable gene (AT2G28810), AGTCGCTGATGTCTTGGGAA and AATCTTCCACAGTCCAGCCA for the highly variable gene (AT1G08930), AACAAGGGCTAACGTGGATG and CTGCTTCTCCTGCTCATTCG for *NRT2.1* (AT1G08090) as well as CCAAGATACAACGCTCAGGC and CAAGGCGTACCCTGCAATCT for AT2G28390, GGCCTTGTATAATCCCTGATGAATAAG and AAAGAGATAACAGGAACGGAAACATAGT for *UBIQUITIN10* (AT4G05320) and finally GCCATCCAAGCTGTTCTCTC and CCCTCGTAGATTGGCACAGT for *ACTIN2* (AT3G18780) the last 3 genes being used as control genes for the normalisation of expression.

### RNA-seq and mapping

RNA quality and integrity were assessed using a Qubit™ Flex Fluorometer (Invitrogen). Generation of libraries and PE150 Illumina sequencing was performed by Novogene (Cambridge, UK) with approximately 20 million reads (of each end) per sample. Quality of reads was assessed using FastQC (www.bioinformatics.babraham.ac.uk/projects/fastqc/) and low quality reads were removed and trimmed using fastp (Chen, 2023). Mapping of the reads on the *Arabidopsis thaliana* genome (TAIR10) was performed with STAR (Dobin et al., 2013) and read count extracted thanks to htseq-count (Anders et al., 2015).

### Data analysis

All data analyses of RNA-seq as well as RTqPCR and luciferase imaging were performed using R (R Core Team, 2021) and scripts are accessible in GitHub (see data availability). Read counts for each gene were normalized by the size of the library (CPM) using scTenifoldNet package (Osorio et al., 2020). Highly variable genes were detected as previously described, as well as the corrected CV for each gene (Cortijo et al., 2019). Hierarchical clustering and heatmap of genes with a corrected CV > 1 was performed on the Zscore (CPM of a gene - mean expression of that gene in samples/ standard deviation of the expression for that gene in samples) using the function hclust on 1-pearson correlation. Gene Ontology enrichment analysis was performed using the shinyGO web application (Ge et al., 2020).

## Data availability

RNA-seq data are on ArrayExpress with the accession number E-MTAB-15039 and will be released once the article is accepted.

Other raw data and script are on GitHub will be made available once the article is accepter.

## Acknowledgments

This work was supported by grants from the I-SITE MUSE (AAP21REC-FRA02, VariPlant) and from the National Agency for Research (ANR-22-CE20-0020, ChromaVari). We acknowledge the Histocytology and Plant Cell Imaging platform (PHIV) and Isotopes Quantifications platform (AQUI) for technical support with luciferase imaging and nitrate influx experiment respectively. We also thank Tou Cheu Xiong for providing the *p35S:LUC* line and Laurine Mancardi for technical help on the Lumalum experiments.

## Authors contribution

SC designed the project, CL, AV and SC implemented the project with help from CF for bioinformatics analysis. HJ performed the Lumalum experiments and MM the transgenerational study. SC, CL, AV and AM wrote the manuscript that was read and corrected by all authors.

## Supplementary figures

**Fig S1.**
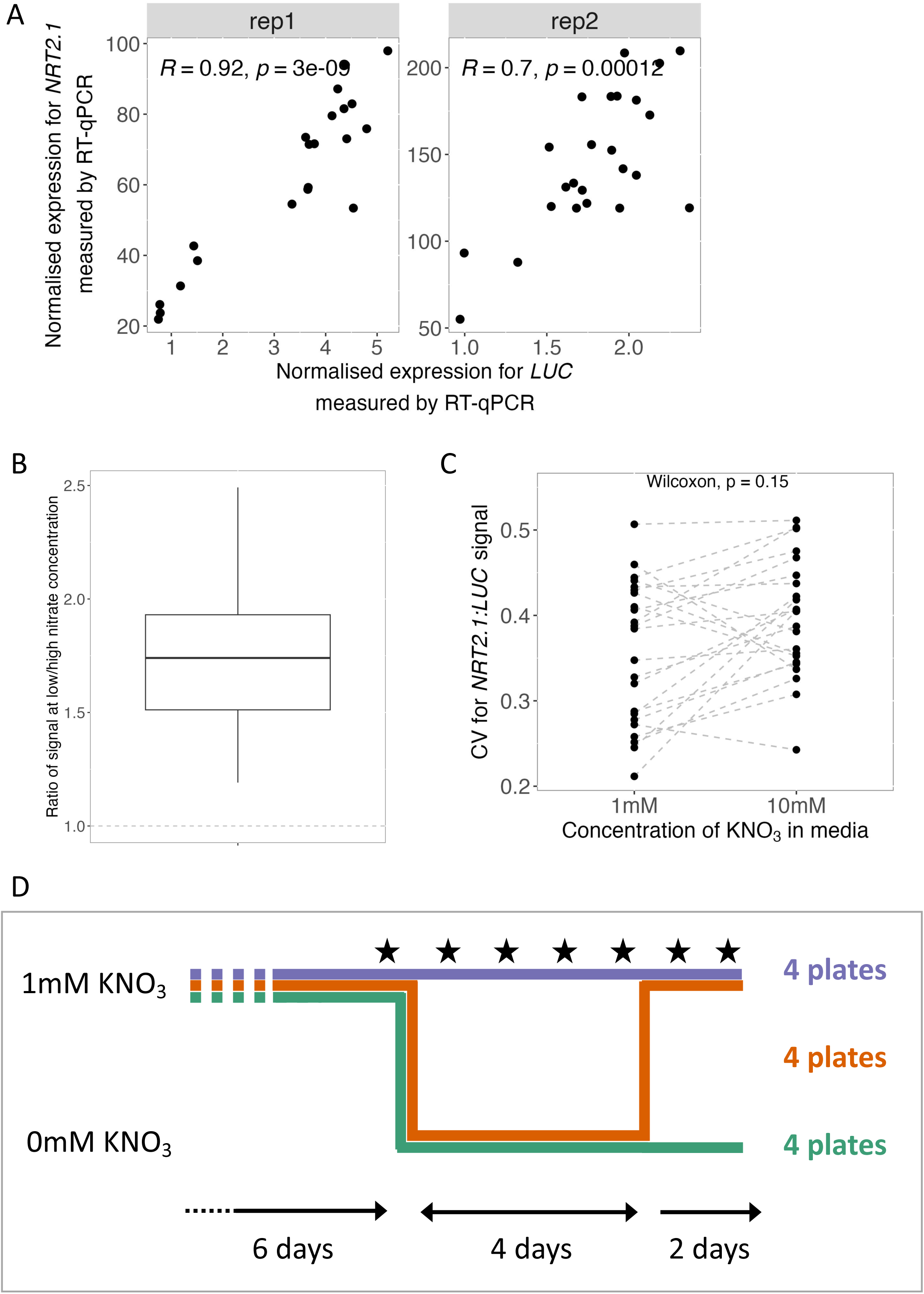
(A) Comparison of the quantity of mRNA measured by RT-qPCR for the *NRT2.1* and luciferase genes in the *pNRT2.1:LUC* reporter line. Each point corresponds to one seedling. The results of a spearman correlation test are included. (B) Distribution of the ratio of the *pNRT2.1:LUC* signal at low (1mM) and high (10mM) nitrate concentration. The dotted line at 1 indicates an absence of difference in signal between low and high nitrate, which is not observed for any of the replicates. (C) Inter-individual transcriptional variability for the *pNRT2.1:LUC* reporter line measured for seedlings grown on a media with low (1mM) of high (10mM) nitrate concentration. Each point represents a replicate with around 25 seedlings on low nitrate and 25 seedlings on high nitrate. Replicates are linked with a grey dotted line. (D) Experimental set-up for the nitrate starvation experiment. Black stars correspond to days when luciferase imaging was performed.

**Fig S2.**
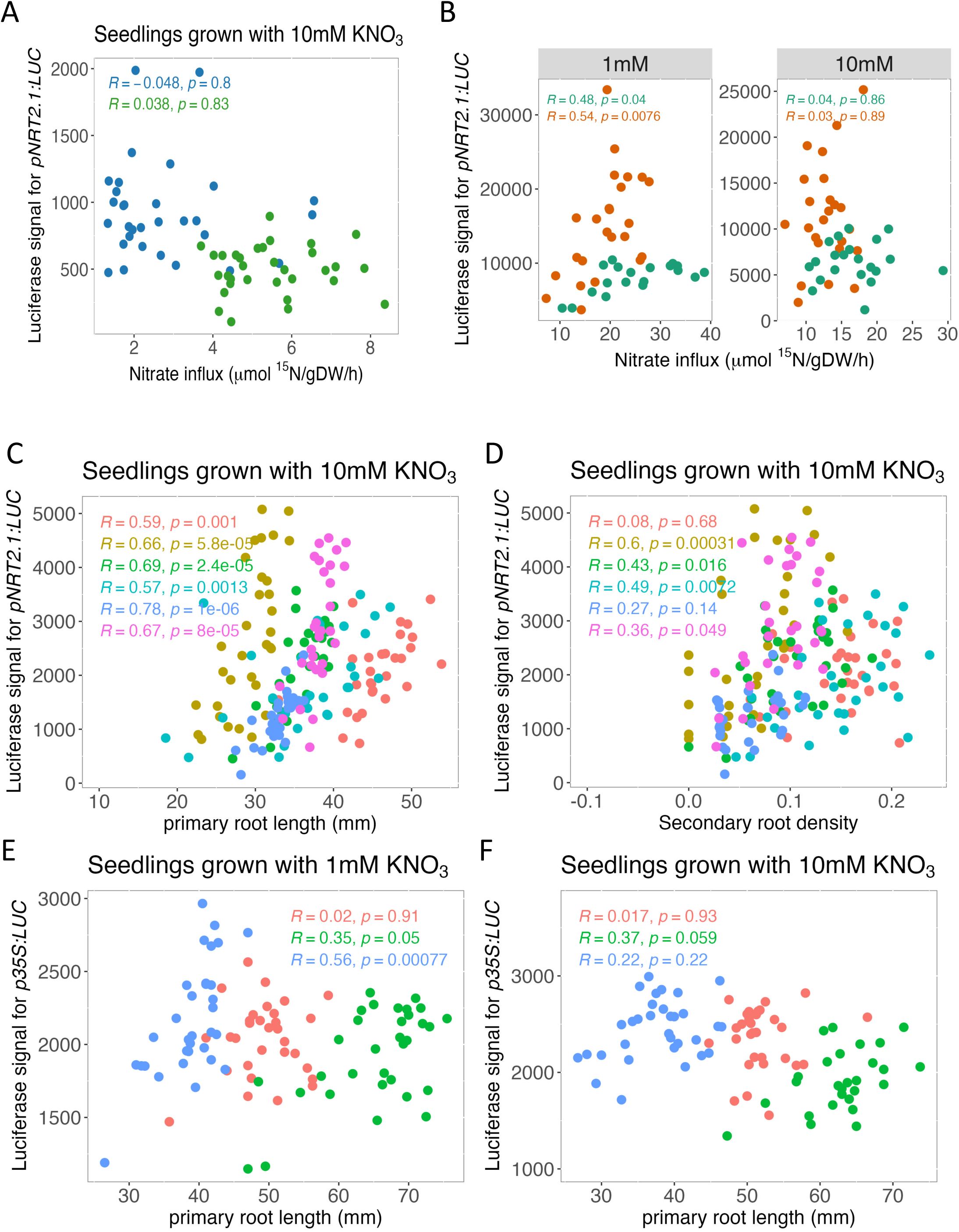
(A) Comparison for plants grown in media with high (10mM) nitrate concentration of the *pNRT2.1:LUC* signal and the nitrate influx of HATS. Each point corresponds to a single seedling and each colour to a replicate. The result of a spearman correlation test for each replicate is included. (B) Comparison of the *pNRT2.1:LUC* signal measured by a second luciferase camera and the nitrate influx of HATS for plants grown with low (left) or high (right) nitrate. Each point corresponds to a single seedling and each colour to a replicate. The result of a spearman correlation test for each replicate is included. (C-D) Comparison for plants grown in media with high (10mM) nitrate concentration of the *pNRT2.1:LUC* signal and (B) the primary root length or (C) the lateral root density. Each point corresponds to a single seedling and each colour to a replicate. The result of a spearman correlation test for each replicate is included. (E-F) Comparison for plants grown in media with low (1mM) nitrate concentration of the *p35S:LUC* signal and (B) the primary root length or (C) the lateral root density. Each point corresponds to a single seedling and each colour to a replicate. The result of a spearman correlation test for each replicate is included.

**Fig S3.**
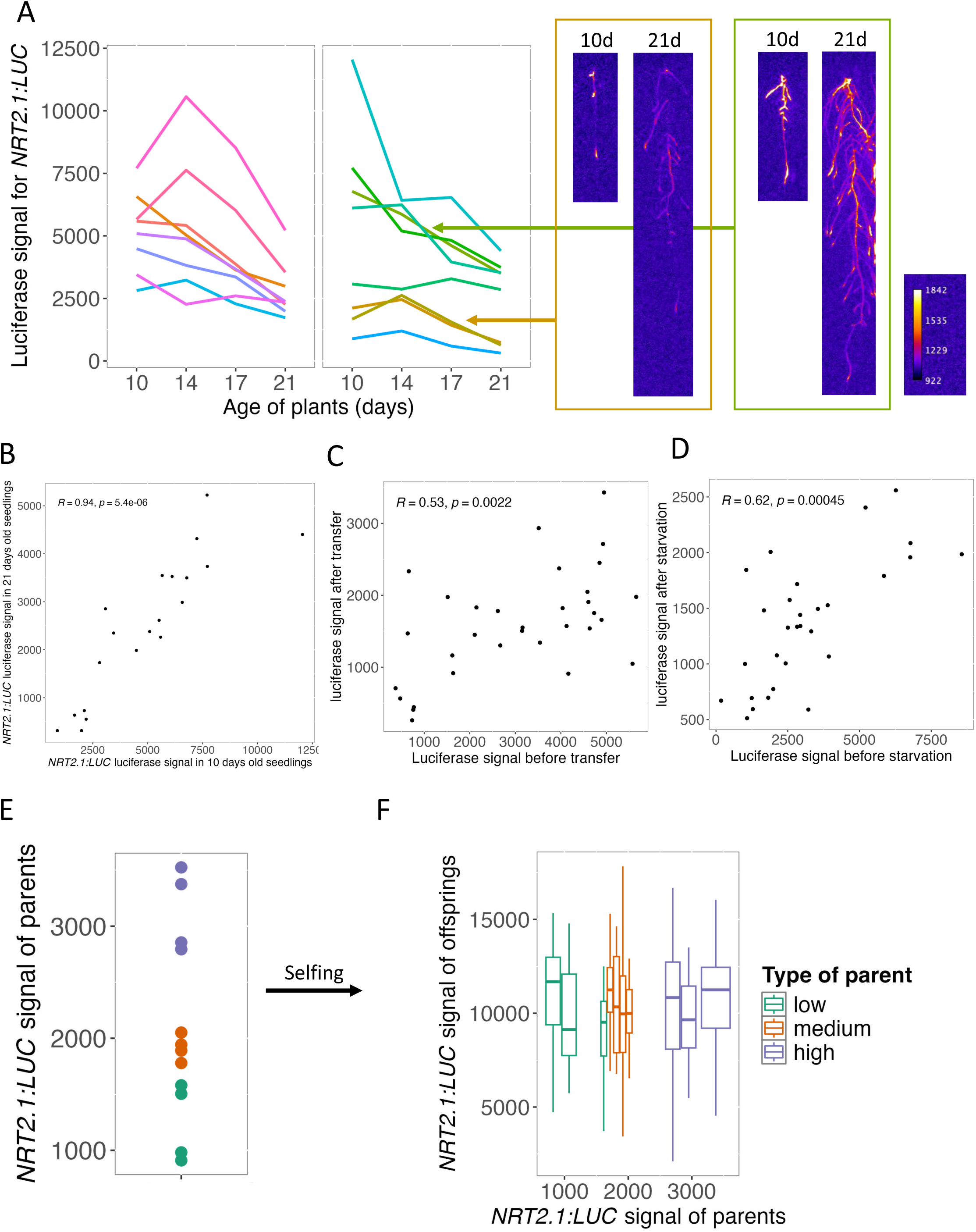
(A) *pNRT2.1:luc* signal over 4 days for two plates. Each line corresponds to a given seedling. (B) Correlation of the *pNRT2.1:LUC* signal in 9-day-old and 12-day-old seedlings. Each point corresponds to a seedling. The result of a spearman correlation test is included. (C) Correlation of the *pNRT2.1:LUC* signal in seedlings before and after a transfer to a new plate with the same concentration of nitrate. Each point corresponds to a seedling. The result of a spearman correlation test is included. (D) Correlation of the *pNRT2.1:LUC* signal in seedlings before and after a nitrate starvation. Each point corresponds to a seedling. The result of a spearman correlation test is included. (E) *pNRT2.1:LUC* signal for 19 seedlings (replicate 2 of Figure 4B). The individuals selected to analyse the signal of their offsprings are shown in colour depending on their category: low expression in green, medium expression in orange and high expression in purple. (F) Distribution of the *pNRT2.1:LUC* signal for populations deriving from self-pollination of the parents selected in (E). Each boxplot corresponds to a population of descendants (at least 24 seedlings), with the colour depending on the category of the parent: low expression in green, medium expression in orange and high expression in purple.

**Fig S4.**
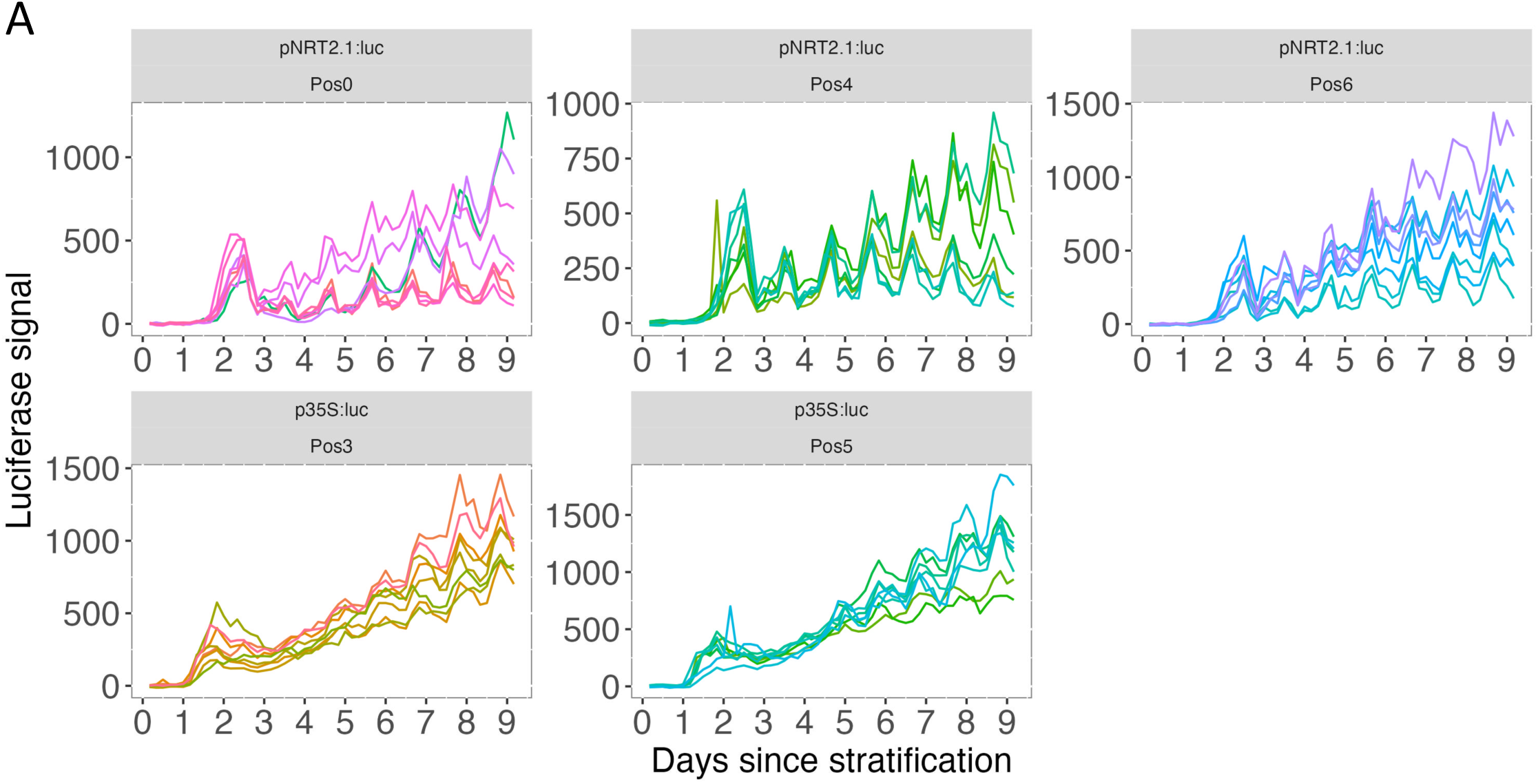
Biological replicate of Figure 5A. *pNRT2.1:LUC* (top) *p35S:LUC* (bottom) *s*ignal over time. Seven to eight seedlings were measured in each plate. Only seedlings that have lateral root at the end of the experiment were measured. Each line represents the signal in a given seedling, measured every 4 hours for 9 days after stratification.

**Fig S5.**
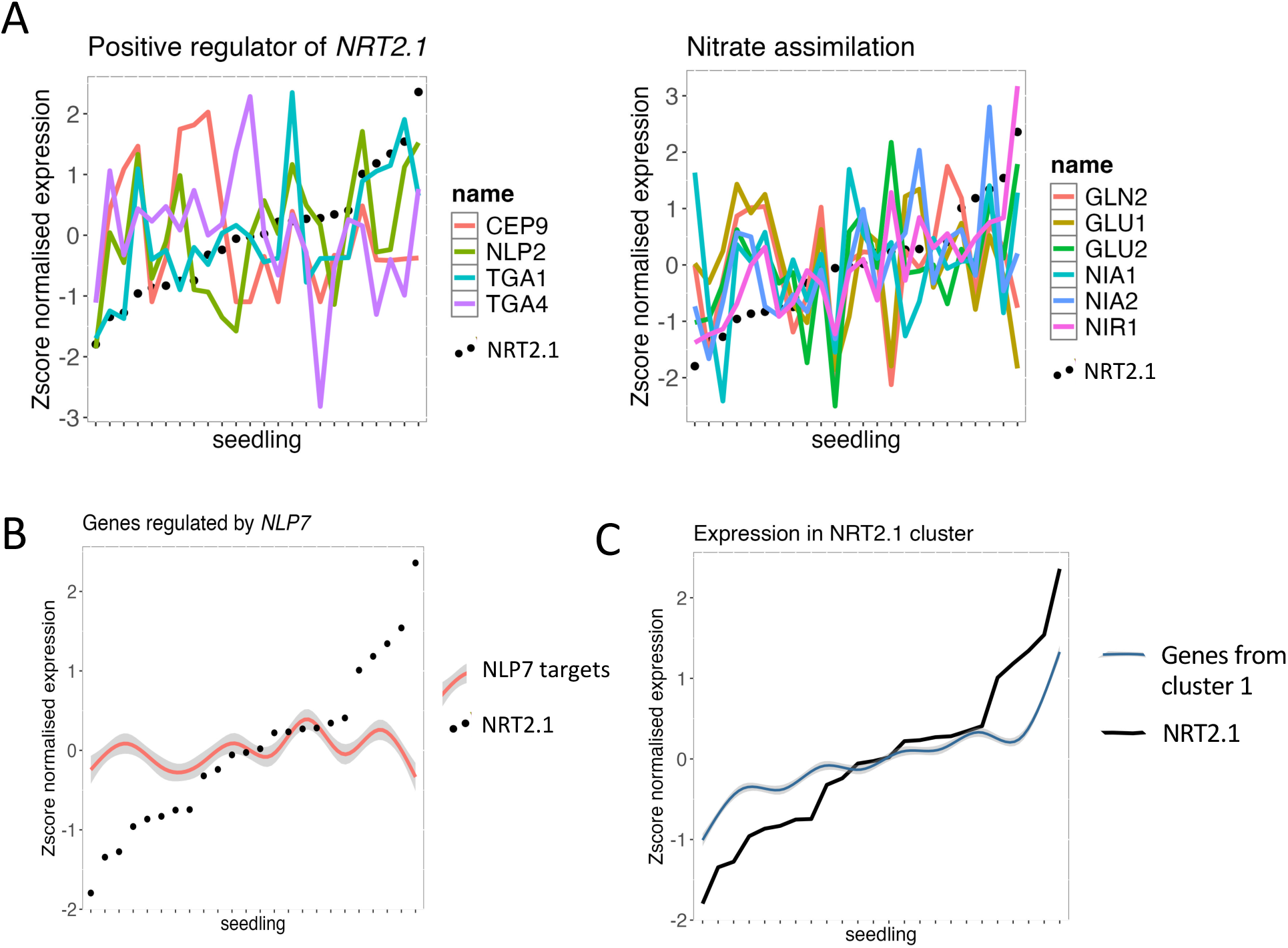
(A) Expression level in the different seedlings of *NRT2.1* (black dots), and of positive regulators of *NRT2.1* expression (left), or genes involved in nitrate assimilation (right). (B) Expression in the different seedlings of *NRT2.1* (black points), and of the average and standard deviation for genes targeted by NLP7 transcription factor. (C) Expression in the different seedlings of *NRT2.1* (black lines), and of the average and standard deviation for the genes in the cluster 1.

## Bibliography

Abley, K., Formosa-Jordan, P., Tavares, H., Chan, E.Y., Afsharinafar, M., Leyser, O. and Locke, J.C. (2021). An ABA-GA bistable switch can account for natural variation in the variability of Arabidopsis seed germination time. Elife. 10:.

Abley, K., Goswami, R. and Locke, J.C.W. (2024). Bet-hedging and variability in plant development: seed germination and beyond. Philos. Trans. R. Soc. Lond. B, Biol. Sci. 379:20230048.

Alamos, S., Reimer, A., Niyogi, K.K. and Garcia, H.G. (2021). Quantitative imaging of RNA polymerase II activity in plants reveals the single-cell basis of tissue-wide transcriptional dynamics. Nat. Plants. 7:1037–1049.

Anders, S., Pyl, P.T. and Huber, W. (2015). HTSeq — a Python framework to work with high-throughput sequencing data. Bioinformatics. 31:166–169.

Araus, V., Vidal, E.A., Puelma, T., Alamos, S., Mieulet, D., Guiderdoni, E. and Gutiérrez, R.A. (2016). Members of BTB gene family of scaffold proteins suppress nitrate uptake and nitrogen use efficiency. Plant Physiol. 171:1523–1532.

Avin-Wittenberg, T., Tzin, V., Angelovici, R., Less, H. and Galili, G. (2012). Deciphering energy-associated gene networks operating in the response of Arabidopsis plants to stress and nutritional cues. The Plant Journal. 70:954–966.

Bellegarde, F., Gojon, A. and Martin, A. (2017). Signals and players in the transcriptional regulation of root responses by local and systemic N signaling in Arabidopsis thaliana. J. Exp. Bot. 68:2553–2565.

Bellegarde, F., Herbert, L., Séré, D., Caillieux, E., Boucherez, J., Fizames, C., Roudier, F., Gojon, A. and Martin, A. (2018). Polycomb Repressive Complex 2 attenuates the very high expression of the Arabidopsis gene NRT2.1. Sci. Rep. 8:7905.

Cain, M.L., Subler, S., Evans, J.P. and Fortin, M.-J. (1999). Sampling spatial and temporal variation in soil nitrogen availability. Oecologia. 118:397–404.

Cerezo, M., Tillard, P., Filleur, S., Muños, S., Daniel-Vedele, F. and Gojon, A. (2001). Major alterations of the regulation of root NO(3)(-) uptake are associated with the mutation of Nrt2.1 and Nrt2.2 genes in Arabidopsis. Plant Physiol. 127:262–271.

Chen, S. (2023). Ultrafast one-pass FASTQ data preprocessing, quality control, and deduplication using fastp. iMeta. 2:e107.

Childs, D.Z., Metcalf, C.J.E. and Rees, M. (2010). Evolutionary bet-hedging in the real world: empirical evidence and challenges revealed by plants. Proc. Biol. Sci. 277:3055–3064.

Cohen, D. (1968). A general model of optimal reproduction in a randomly varying environment. J Ecol. 56:219.

Cortijo, S., Aydin, Z., Ahnert, S. and Locke, J.C. (2019). Widespread inter-individual gene expression variability in Arabidopsis thaliana. Mol. Syst. Biol. 15:e8591.

Cortijo, S. and Locke, J.C.W. (2020). Does gene expression noise play a functional role in plants? Trends Plant Sci. 25:1041–1051.

Dey, S.S., Foley, J.E., Limsirichai, P., Schaffer, D.V. and Arkin, A.P. (2015). Orthogonal control of expression mean and variance by epigenetic features at different genomic loci. Mol. Syst. Biol. 11:806.

Dobin, A., Davis, C.A., Schlesinger, F., Drenkow, J., Zaleski, C., Jha, S., Batut, P., Chaisson, M. and Gingeras, T.R. (2013). STAR: ultrafast universal RNA-seq aligner. Bioinformatics. 29:15–21.

Dong, P. and Liu, Z. (2017). Shaping development by stochasticity and dynamics in gene regulation. Open Biol. 7:.

Dorbe, M.-F., Truong, H.-N., Crété, P. and Daniel-Vedele, F. (1998). Deletion analysis of the tobacco Nii1 promoter in Arabidopsis thaliana. Plant Sci. 139:71–82.

Filleur, S., Dorbe, M.F., Cerezo, M., Orsel, M., Granier, F., Gojon, A. and Daniel-Vedele, F. (2001). An arabidopsis T-DNA mutant affected in Nrt2 genes is impaired in nitrate uptake. FEBS Lett. 489:220–224.

Forde, B.G. (2009). Is it good noise? The role of developmental instability in the shaping of a root system. J. Exp. Bot. 60:3989–4002.

Gasch, A.P. et al. (2017). Single-cell RNA sequencing reveals intrinsic and extrinsic regulatory heterogeneity in yeast responding to stress. PLoS Biol. 15:e2004050.

Ge, S.X., Jung, D. and Yao, R. (2020). ShinyGO: a graphical gene-set enrichment tool for animals and plants. Bioinformatics. 36:2628–2629.

Girin, T., El-Kafafi, E.-S., Widiez, T., Erban, A., Hubberten, H.-M., Kopka, J., Hoefgen, R., Gojon, A. and Lepetit, M. (2010). Identification of Arabidopsis mutants impaired in the systemic regulation of root nitrate uptake by the nitrogen status of the plant. Plant Physiol. 153:1250–1260.

Graham, I.A. (2008). Seed storage oil mobilization. Annu. Rev. Plant Biol. 59:115–142.

Grimbergen, A.J., Siebring, J., Solopova, A. and Kuipers, O.P. (2015). Microbial bet-hedging: the power of being different. Curr. Opin. Microbiol. 25:67–72.

Hall, M.C., Dworkin, I., Ungerer, M.C. and Purugganan, M. (2007). Genetics of microenvironmental canalization in Arabidopsis thaliana. Proc. Natl. Acad. Sci. USA. 104:13717–13722.

Hanchi, M. et al. (2018). The Phosphate Fast-Responsive Genes PECP1 and PPsPase1 Affect Phosphocholine and Phosphoethanolamine Content. Plant Physiol. 176:2943–2962.

Jimenez-Gomez, J.M., Corwin, J.A., Joseph, B., Maloof, J.N. and Kliebenstein, D.J. (2011). Genomic analysis of QTLs and genes altering natural variation in stochastic noise. PLoS Genet. 7:e1002295.

Joseph, B., Corwin, J.A. and Kliebenstein, D.J. (2015). Genetic variation in the nuclear and organellar genomes modulates stochastic variation in the metabolome, growth, and defense. PLoS Genet. 11:e1004779.

Kiba, T. et al. (2018). Repression of Nitrogen Starvation Responses by Members of the Arabidopsis GARP-Type Transcription Factor NIGT1/HRS1 Subfamily. Plant Cell. 30:925–945.

Kiba, T. et al. (2012). The Arabidopsis nitrate transporter NRT2.4 plays a double role in roots and shoots of nitrogen-starved plants. Plant Cell. 24:245–258.

Kircher, S. and Schopfer, P. (2012). Photosynthetic sucrose acts as cotyledon-derived long-distance signal to control root growth during early seedling development in Arabidopsis. Proc. Natl. Acad. Sci. USA. 109:11217–11221.

Kitazawa, M.S. (2021). Developmental stochasticity and variation in floral phyllotaxis. J Plant Res. 134:403–416.

Kusuma, S.H., Kakizuka, T., Hattori, M. and Nagai, T. (2024). Autonomous multicolor bioluminescence imaging in bacteria, mammalian, and plant hosts. Proc. Natl. Acad. Sci. USA. 121:e2406358121.

Laugier, E., Bouguyon, E., Mauriès, A., Tillard, P., Gojon, A. and Lejay, L. (2012). Regulation of high-affinity nitrate uptake in roots of Arabidopsis depends predominantly on posttranscriptional control of the NRT2.1/NAR2.1 transport system. Plant Physiol. 158:1067–1078.

Lejay, L., Tillard, P., Lepetit, M., Olive, F. d, Filleur, S., Daniel-Vedele, F. and Gojon, A. (1999). Molecular and functional regulation of two NO3-uptake systems by N- and C-status of Arabidopsis plants. The Plant Journal. 18:509–519.

Lezhneva, L., Kiba, T., Feria-Bourrellier, A.-B., Lafouge, F., Boutet-Mercey, S., Zoufan, P., Sakakibara, H., Daniel-Vedele, F. and Krapp, A. (2014). The Arabidopsis nitrate transporter NRT2.5 plays a role in nitrate acquisition and remobilization in nitrogen-starved plants. The Plant Journal. 80:230–241.

Lin, L., Cao, G., Zhang, F., Ke, X., Li, Y., Xu, X., Li, Q., Guo, X., Fan, B. and Du, Y. (2019). Spatial and Temporal Variations in Available Soil Nitrogen—A Case Study in Kobresia Alpine Meadow in the Qinghai-Tibetan Plateau, China. GEP. 07:177–189.

Little, D.Y., Rao, H., Oliva, S., Daniel-Vedele, F., Krapp, A. and Malamy, J.E. (2005). The putative high-affinity nitrate transporter NRT2.1 represses lateral root initiation in response to nutritional cues. Proc. Natl. Acad. Sci. USA. 102:13693–13698.

Marchive, C., Roudier, F., Castaings, L., Bréhaut, V., Blondet, E., Colot, V., Meyer, C. and Krapp, A. (2013). Nuclear retention of the transcription factor NLP7 orchestrates the early response to nitrate in plants. Nat. Commun. 4:1713.

Martin, T., Oswald, O. and Graham, I.A. (2002). Arabidopsis seedling growth, storage lipid mobilization, and photosynthetic gene expression are regulated by carbon:nitrogen availability. Plant Physiol. 128:472–481.

Medici, A. and Krouk, G. (2014). The primary nitrate response: a multifaceted signalling pathway. J. Exp. Bot. 65:5567–5576.

Mitchell, J., Johnston, I.G. and Bassel, G.W. (2017). Variability in seeds: biological, ecological, and agricultural implications. J. Exp. Bot. 68:809–817.

Morawska, L.P., Hernandez-Valdes, J.A. and Kuipers, O.P. (2022). Diversity of bet-hedging strategies in microbial communities-Recent cases and insights. WIREs Mechanisms of Disease. 14:e1544.

Nazoa, P., Vidmar, J.J., Tranbarger, T.J., Mouline, K., Damiani, I., Tillard, P., Zhuo, D., Glass, A.D.M. and Touraine, B. (2003). Regulation of the nitrate transporter gene AtNRT2.1 in Arabidopsis thaliana: responses to nitrate, amino acids and developmental stage. Plant Mol. Biol. 52:689–703.

O’Brien, J.A., Vega, A., Bouguyon, E., Krouk, G., Gojon, A., Coruzzi, G. and Gutiérrez, R.A. (2016). Nitrate transport, sensing, and responses in plants. Mol. Plant. 9:837–856.

Ohkubo, Y., Kuwata, K. and Matsubayashi, Y. (2021). A type 2C protein phosphatase activates high-affinity nitrate uptake by dephosphorylating NRT2.1. Nat. Plants. 7:310–316.

Orsel, M., Chopin, F., Leleu, O., Smith, S.J., Krapp, A., Daniel-Vedele, F. and Miller, A.J. (2007). Nitrate signaling and the two component high affinity uptake system in Arabidopsis. Plant Signal. Behav. 2:260–262.

Osorio, D., Zhong, Y., Li, G., Huang, J.Z. and Cai, J.J. (2020). scTenifoldNet: A Machine Learning Workflow for Constructing and Comparing Transcriptome-wide Gene Regulatory Networks from Single-Cell Data. Patterns (N Y). 1:100139.

R Core Team (2021). R: A language and environment for statistical. R Foundation for Statistical Computing, Vienna, Austria.

Remans, T., Nacry, P., Pervent, M., Girin, T., Tillard, P., Lepetit, M. and Gojon, A. (2006). A central role for the nitrate transporter NRT2.1 in the integrated morphological and physiological responses of the root system to nitrogen limitation in Arabidopsis. Plant Physiol. 140:909–921.

Rubin, G., Tohge, T., Matsuda, F., Saito, K. and Scheible, W.-R. (2009). Members of the LBD family of transcription factors repress anthocyanin synthesis and affect additional nitrogen responses in Arabidopsis. Plant Cell. 21:3567–3584.

Ruffel, S. et al. (2019). Genome-wide Analysis In Response to N and C Identifies New Regulators for root AtNRT2 Transporters. BioRxiv.

Samo, N. et al. (2024). PRC2 facilitates the transition from heterotrophy to photoautotrophy during seedling emergence. BioRxiv.

Schindelin, J. et al. (2012). Fiji: an open-source platform for biological-image analysis. Nat. Methods. 9:676–682.

Shahandeh, H., Wright, A.L., Hons, F.M. and Lascano, R.J. (2005). Spatial and temporal variation of soil nitrogen parameters related to soil texture and corn yield. Agron J. 97:772–782.

Shakhova, E.S. et al. (2024). An improved pathway for autonomous bioluminescence imaging in eukaryotes. Nat. Methods. 21:406–410.

Sharon, E., Dijk, D. van, Kalma, Y., Keren, L., Manor, O., Yakhini, Z. and Segal, E. (2014). Probing the effect of promoters on noise in gene expression using thousands of designed sequences. Genome Res. 24:1698–1706.

Shen, X., Pettersson, M., Rönnegård, L. and Carlborg, Ö. (2012). Inheritance beyond plain heritability: variance-controlling genes in Arabidopsis thaliana. PLoS Genet. 8:e1002839.

Silva, A.T., Ribone, P.A., Chan, R.L., Ligterink, W. and Hilhorst, H.W.M. (2016). A Predictive Coexpression Network Identifies Novel Genes Controlling the Seed-to-Seedling Phase Transition in Arabidopsis thaliana. Plant Physiol. 170:2218–2231.

Simon, L. and Probst, A.V. (2024). Maintenance and dynamic reprogramming of chromatin organization during development. The Plant Journal. 118:657–670.

Tantale, K. et al. (2016). A single-molecule view of transcription reveals convoys of RNA polymerases and multi-scale bursting. Nat. Commun. 7:12248.

Tan-Wilson, A.L. and Wilson, K.A. (2012). Mobilization of seed protein reserves. Physiol. Plant. 145:140–153.

Tirosh, I. and Barkai, N. (2008). Two strategies for gene regulation by promoter nucleosomes. Genome Res. 18:1084–1091.

Urchueguía, A., Galbusera, L., Chauvin, D., Bellement, G., Julou, T. and Nimwegen, E. van (2021). Genome-wide gene expression noise in Escherichia coli is condition-dependent and determined by propagation of noise through the regulatory network. PLoS Biol. 19:e3001491.

Wang, Y., Yuan, Z., Wang, J., Xiao, H., Wan, L., Li, L., Guo, Y., Gong, Z., Friml, J. and Zhang, J. (2023). The nitrate transporter NRT2.1 directly antagonizes PIN7-mediated auxin transport for root growth adaptation. Proc. Natl. Acad. Sci. USA. 120:e2221313120.

Wang, Y.-Y., Hsu, P.-K. and Tsay, Y.-F. (2012). Uptake, allocation and signaling of nitrate. Trends Plant Sci. 17:458–467.

Wirth, J., Chopin, F., Santoni, V., Viennois, G., Tillard, P., Krapp, A., Lejay, L., Daniel-Vedele, F. and Gojon, A. (2007). Regulation of root nitrate uptake at the NRT2.1 protein level in Arabidopsis thaliana. J. Biol. Chem. 282:23541–23552.

Zhuo, D., Okamoto, M., Vidmar, J.J. and Glass, A.D. (1999). Regulation of a putative high-affinity nitrate transporter (Nrt2;1At) in roots of Arabidopsis thaliana. The Plant Journal. 17:563–568.

